# Constitutive protein degradation induces acute cell death via proteolysis products

**DOI:** 10.1101/2023.02.06.527237

**Authors:** Si-Han Chen, Sumit Prakash, Elizabeth Helgason, Caroline L. Gilchrist, Lillian R. Kenner, Rajini Srinivasan, Tim Sterne-Weiler, Marc Hafner, Robert Piskol, Erin C. Dueber, Habib Hamidi, Nicholas Endres, Xin Ye, Wayne J. Fairbrother, Ingrid E. Wertz

**Author notes:** Current address: Lyterian Therapeutics, 1 Letterman Drive, Building D, Suite DM-900, San Francisco, CA 94129, USA. Current address: Bristol Myers Squibb, 10300 Campus Point Dr Suite 100, San Diego, CA 92121, USA.

## Abstract

Modulation of proteolysis is an emerging therapeutic mainstay. The clinical success of thalidomide and analogs has inspired development of rationally-designed therapeutics that repurpose endogenous degradation machinery to target pathogenic proteins. However, it is unknown whether target removal is the critical effect that drives degrader-induced efficacy. Here we report that proteasome-generated peptides actively initiate degrader-induced cell death. Utilizing BET family degraders as exemplars, we find that induced proteasomal degradation of the BRD4-long isoform (BRD4-L) generates neo-amino-terminal peptides that neutralize Inhibitor of Apoptosis (IAP) proteins to precipitate cell death. Depletion of BRD4-L paradoxically suppresses caspase activation induced by numerous BET degraders. An unbiased screen revealed that other degrader compounds, including clinical CELMoDs, rely on the same mechanism to potentiate caspase activation and apoptosis. Finally, in the context of constitutive immunoglobulin proteostasis within multiple myeloma cells, we report that therapeutic proteasomal protease inhibition alters the peptide repertoire to neutralize IAPs, thus contributing to the clinical efficacy of bortezomib. Together, these findings clarify the counterintuitive clinical benefit achieved by combining thalidomide analogs with proteasome inhibitors. Our study reveals a previously unrealized pro-apoptotic function of the peptides generated by a variety of proteolysis-modulating compounds, that provide design considerations to maximize therapeutic benefit.

## Main

The ubiquitin-proteasome system governs protein homeostasis by mediating ubiquitination of unwanted proteins. The resulting ubiquitin conjugates then direct the proteasomal degradation of the targeted substrates. Certain therapeutic and preclinical compounds have been serendipitously discovered to function as molecular glues, that induce proximity of ubiquitin ligases (E3) with neosubstrates, resulting in ubiquitination and subsequent degradation of the recruited substrates. For example, thalidomide and related derivatives are molecular glue degraders that achieve their therapeutic properties by targeting neosubstrates that modulate the immune response and drive tumorigenesis for degradation^1, 2^. Furthermore, thalidomide analogs can be incorporated into heterobifunctional protein degraders to achieve potent target degradation^3, 4^, and are being evaluated in the clinic.

The strategy of designing compounds that promote Targeted Protein Degradation (TPD) of therapeutically relevant targets has advanced rapidly^5^. Target removal by TPD promises a complete loss of function, provided that degradation kinetics exceeds the rate of target protein resynthesis. However, it is unknown whether there are additional on-target effects beyond target removal that contribute to degrader-induced therapeutic efficacy. TPD is one component of the broader therapeutic strategy to modulate proteolysis, that also includes proteasome protease inhibition. Paradoxically, proteasomal protease inhibitors such as bortezomib used in combination with thalidomide analogs are a therapeutic mainstay for certain hematologic malignancies. How these two types of functionally opposing small molecules synergize to drive clinical efficacy remains unknown^6^.

Here, we use systematic and hypothesis-driven approaches to reveal the molecular mechanisms that explain how proteolysis-modulating compounds constitutively produce proteasome-generated peptides, that engage Inhibitor of Apoptosis (IAP) proteins to precipitate cell death and drive therapeutic efficacy. We use well-characterized heterobifunctional degraders of the Bromodomain and Extra-Terminal (BET) family of proteins as initial exemplars to elucidate the cellular machinery and show that therapeutic protein degraders, including thalidomide analog Cereblon E3 Ligase Modulators (CELMoDs) and proteasome protease inhibitors, also function via the same mechanism. In so doing, we outline a screening strategy for proteolysis-regulating compounds that induce this cellular consequence. Our collective studies highlight a previously unrecognized mechanism that drives cell death in response to a variety of proteolysis-modulating small molecules. These findings serve as the basis for designing compounds that intentionally exploit or uncouple this cytotoxic effect, in order to maximize degrader-induced efficacy and minimize toxicity for optimal therapeutic benefit.

## Cytotoxicity of BET degradation is not recapitulated by BET inhibition or depletion

Degradation of the BET family of proteins (including BRD4, BRD3, and BRD2) is a well-established example in which degraders consistently outperform their inhibitor counterparts, despite having less favorable drug-like properties^7^. Specifically, BET degraders (BETd) promote caspase activation in both hematological and solid tumor models^4, 8^. The enhanced cytotoxicity achieved by BETd over BET inhibitors is assumed to result from target removal^9, 10^. As such, we used a panel of well-defined BETd for our initial studies (Supplementary Fig. 1).

To quantitatively assess cytostatic versus cytotoxic responses of BET inhibition and degradation, we took advantage of growth rate-based (GR) drug sensitivity metrics. GRmax mirrors conventional Emax in representing drug efficacy and is also scaled by assay duration, drug treatment time and cell doubling time^11^. Positive GRmax values between 1 and 0 indicate partial or complete cytostasis, whereas negative GRmax values down to -1 are indicative of cytotoxicity. First, we assessed cellular responses to the VHL-recruiting BETd compound MZ1^12^ and its inactive epimer cis-MZ1 as the orthogonal BETi control in a panel of cell line models, encompassing AR-positive prostate cancer^13^, ER-positive breast cancer^14^, and hematologic malignancies^15^. MZ1 is overwhelmingly cytotoxic with 17 out of 18 cell lines showing negative GRmax values; by contrast, cis-MZ1 only shows cytotoxicity in 5 cell lines (Fig. 1a). Comparison with the parental BETi ligand JQ1 in an expanded panel of 192 cell lines further confirmed the enhanced cytotoxic response of BETd over BETi (Extended Data Fig. 1).

**Fig. 1.**
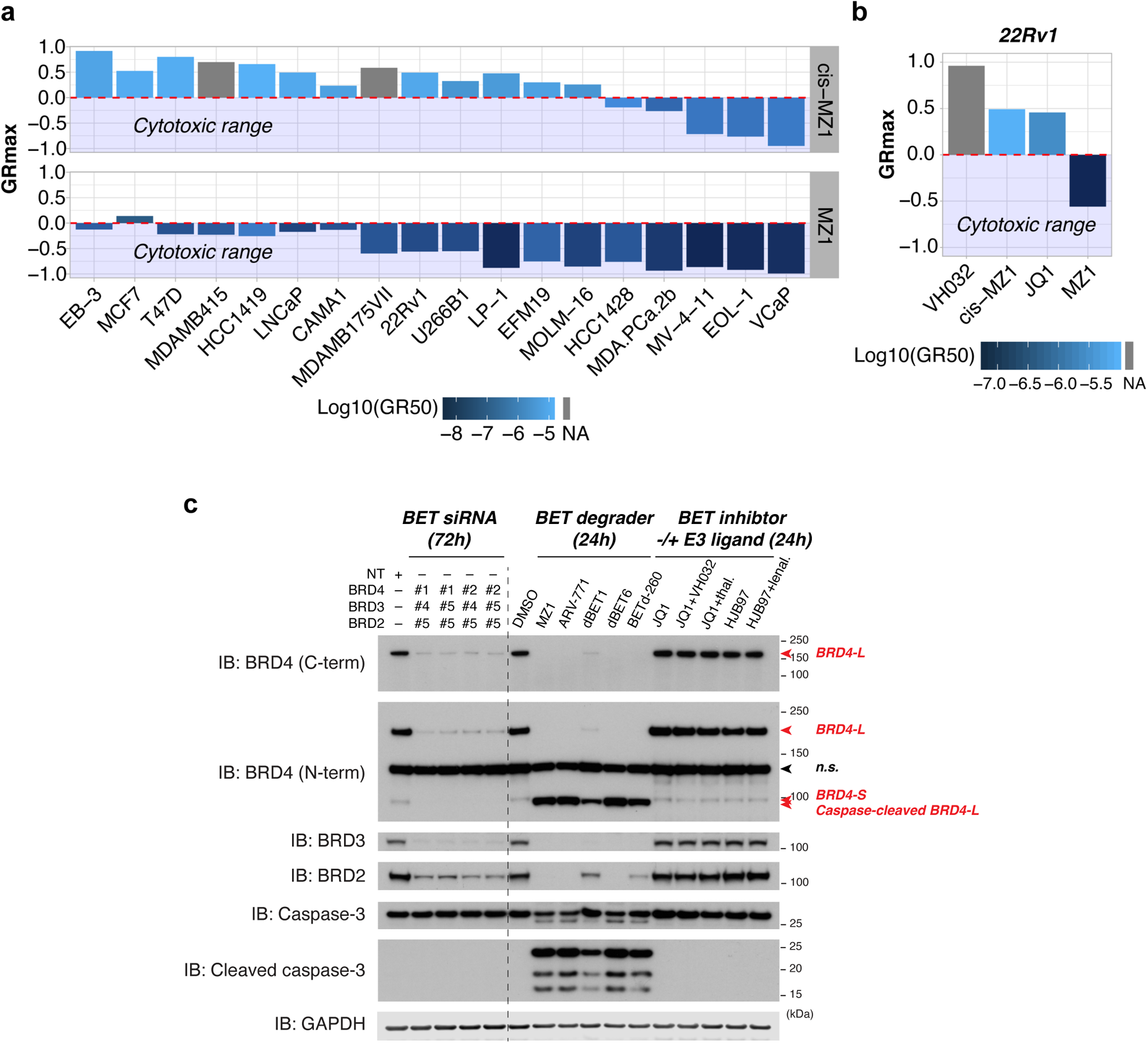
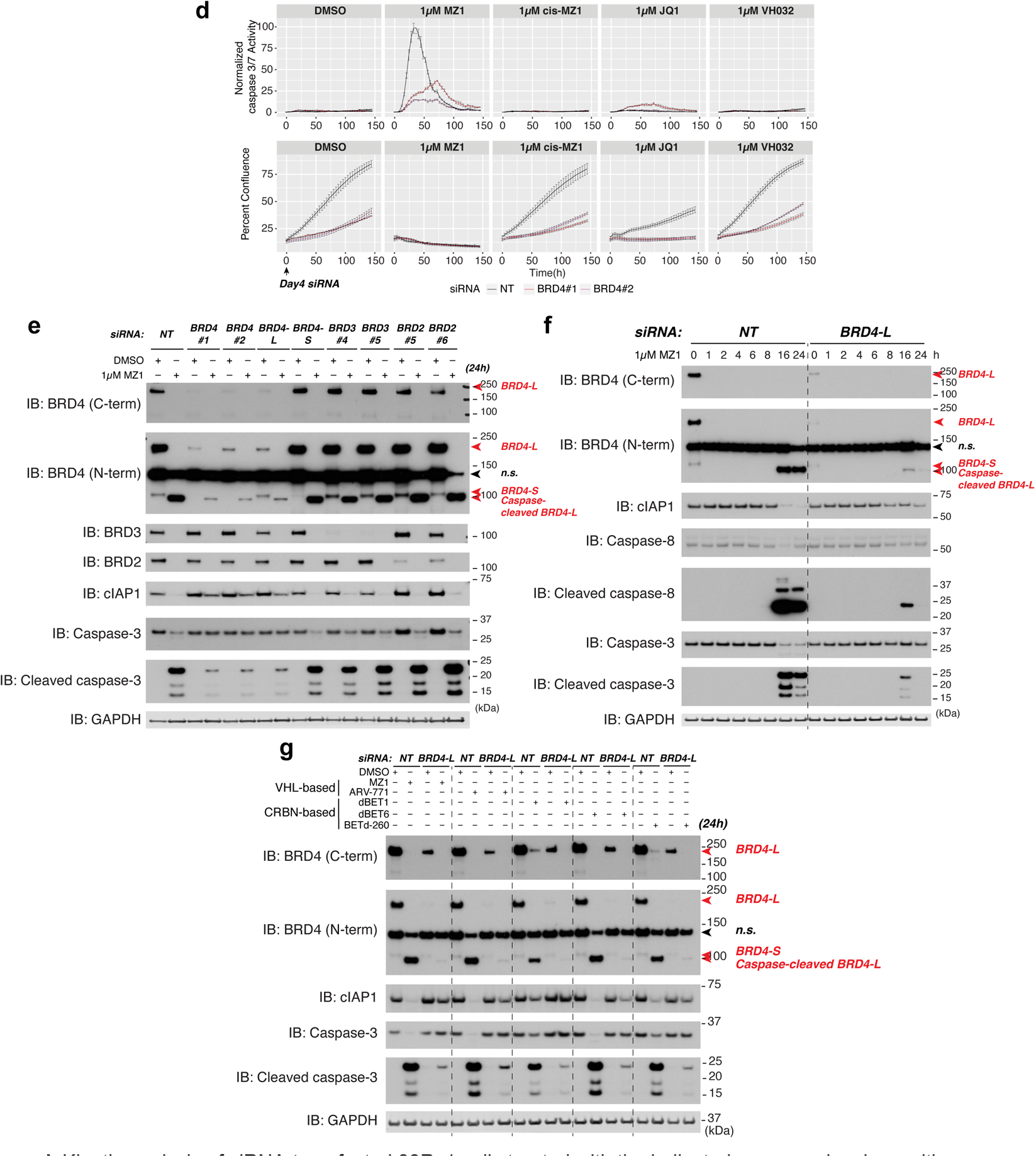
| BET degraders activate caspases via a BRD4-L-dependent manner. **a**, Growth-rate (GR) drug sensitivity metrics of human cancer cell lines in response to the BET degrader MZ1 and the inactive epimer cis-MZ1, calculated based on dose-response curves from triplicate experiments. GRmax less than zero denotes cytotoxic range (efficacy). Log-transformed GR50 represents drug potency. NA denotes unavailable GR50 due to insensitivity to the treatments. **b**, GR metrics of the 22Rv1 cell line in response to MZ1, the inhibitor controls JQ1 and cis-MZ1, and the E3 ligand VH032. **c**, Immunoblot of the indicated proteins from 22Rv1 cells treated with BET siRNA, degrader or inhibitor in combination with the corresponding E3 ligand. C-term and N-term denote two separate BRD4 antibodies that recognize the amino acids 1312–1362 and 150–250 of BRD4, respectively. BET inhibitor -/+ E3 ligand: JQ1 and the paired JQ1 + VH032 are controls for MZ1 and ARV-771; JQ1 and JQ1 + thal.(thalidomide) for dBET1 and dBET6; HJB97 and HJB97 + lenal. (lenalidomide) for BETd-260 (also see Supplementary Fig. 1 for tabulated BETd molecules and the corresponding parental ligands). NT, non-targeting; n.s., non-specific bands. **d**, Kinetic analysis of siRNA-transfected 22Rv1 cells treated with the indicated compounds, along with Caspase-3/7 Green Detection Reagent (Invitrogen). Time zero denotes 4 days after siRNA transfection and the beginning of drug treatments. Data are presented as mean ± se; n = 3. **e**, siRNA-treated 22Rv1 cells were re-plated for DMSO or MZ1 treatment and collected for immunoblot by the indicated time point. **f**, Time course analysis of MZ1 treatments in siRNA-treated 22Rv1 cells. **g**, siRNA-treated 22Rv1 cells were re-plated for the treatment of DMSO or the indicated BETd compound. Unless stated otherwise in Reporting Summary, data represent two to four independent experiments. Source data are provided in Supplementary Information.

BETd cytotoxicity is reported to arise from acute loss of BRD4 and subsequent global transcriptional suppression, that predicts BETd will target BETi-sensitive cells as well as broader subpopulations within a given cell line^9, 10^. To map the cell line subpopulations that are depleted by or survive drug treatments to their pre-treatment transcriptomic profiles, we performed TraCe-seq (tracking differential clonal response by single-cell RNA sequencing)^16^ in the 22Rv1 prostate carcinoma cell line. While MZ1 showed pronounced cytotoxicity relative to cis-MZ1 and JQ1 (Fig. 1b), TraCe-seq surprisingly revealed little or no overlap between MZ1- and JQ1-resistant subpopulations as shown by their distinct clonal barcodes and pre-treatment transcriptomes, indicating that MZ1 and JQ1 target different 22Rv1 subpopulations (Extended Data Fig. 2).

Next, we asked whether triple knockdown of BET family proteins, BRD4/3/2, via RNA interference (RNAi) recapitulates BETd cytotoxicity in 22Rv1. Strikingly, BET depletion by siRNA did not elicit caspase activation, despite having a knockdown efficiency comparable to dBET1 (Fig. 1c). By contrast, all BETd compounds evaluated induced caspase-3 cleavage as well as a caspase-cleaved BRD4-L product, which were not observed in cells treated with BETi and BETi in combination with the relevant E3 ligands (Fig. 1c; Supplementary Discussion). Collectively, these data strongly suggest that BET depletion does not fully explain BETd cytotoxicity.

## Caspase activation by BET degraders requires BRD4-L

These observations prompted us to investigate how BET depletion and BET degradation impact cell confluence and caspase activity. To this end, we exposed non-targeting (NT) control and BRD4 siRNA-transfected cells at the same seeding density to MZ1 or control compounds, and monitored the treated cells over time. BRD4 depletion alone slowed cell proliferation and yet did not activate caspases in 22Rv1 cells throughout the course of the experiment, despite sustained knockdown (Fig. 1d, Extended Data Fig. 3a). By contrast, MZ1 induced acute and potent caspase activation, which was unexpectedly suppressed by BRD4 knockdown, indicating that the presence of BRD4 and its degradation per se were required for the observed caspase activity. The control condition, combining BRD4 depletion and JQ1 treatment, collectively inhibited cell growth but induced minimal caspase activation. To further understand this rescue effect, we tracked and compared the confluence of 22Rv1 cells that were exposed to BETd or non-BETd cytotoxic agents in the presence or the absence of BRD4 siRNA. BRD4 depletion by siRNA suppressed cell death between 10 to 40 hours after BETd treatments, but did not affect treatments with non-BETd cytotoxic agents, further implicating BETd-induced BRD4 proteolysis, and not BET protein removal, in BETd-induced caspase activation (Extended Data Fig. 3b).

We next asked whether the degradation of BRD4 was solely responsible for BETd-induced caspase activation, given the collateral degradation of BRD2 and BRD3 by MZ1. Thus, we transfected 22Rv1 cells with BRD2, BRD3, pan-BRD4 or isoform-specific BRD4 siRNAs, followed by MZ1 treatment. Only pan-BRD4 and BRD4-L isoform-specific siRNAs suppressed the caspase-3 activation, indicating MZ1 induces caspase activation and cell killing specifically through BRD4-L (Fig. 1e, Extended Data Fig. 3c). By contrast, BRD4-L was dispensable for caspase activation by non-BETd cytotoxic agents, suggesting that BRD4-L is specifically required by BETd for caspase activation (Extended Data Fig. 3d). Lastly, the rescue of BETd-induced caspase activation by BRD4 depletion was recapitulated by different BETd molecules and across other cell lines (Extended Data Fig. 4). These results indicate that BETd-induced caspase activation is a gain-of-function effect that requires the presence of BRD4-L. BRD4 knockdown alone promotes cytostasis after 50 hours (Fig. 1d); notably, knockdown of BRD4-L protein expression specifically abrogates BETd-induced BRD4-L proteolysis and acute (10–50 hour) caspase activation. It is this acute BETd-initiated caspase activation that likely drives the enhanced cytotoxicity induced by BETd relative to the cytostasis induced by BETi (Fig. 1a, Extended Data Fig. 1).

Next, we explored how BETd-induced BRD4-L degradation promotes caspase activation. A rich body of literature has shown that caspase activation is associated with the exposure of neo-amino termini resulting from autocatalytic cleavage of caspases or from the release of Smac/DIABLO and Smac-like proteins from the mitochondria^17–19^. To avoid inadvertent activation of caspases, the IAP (inhibitors of apoptosis) family of proteins recognize caspase neo-amino termini that contain IAP-binding motifs (IBMs), resulting in direct inhibition or targeting of caspases for proteasomal degradation via the E3 activity of IAP proteins^20, 21^. However, when the cytosolic accumulation of IBMs, such as Smac/DIABLO, overcomes the apoptotic threshold maintained by IAP proteins, caspase activation ensues^22^. Thus, we hypothesized that BETd activates caspases by producing IBM peptides via BRD4-L proteasomal degradation. To test this, we tracked cIAP1 levels, because cIAP1 is autoubiquitinated and degraded by the proteasome upon IBM engagement^23–25^. Indeed, cIAP1 levels were reduced by MZ1 treatment; importantly, prior BRD4-L depletion by siRNA prevented MZ1-induced loss of cIAP1 (Fig. 1e). Genetic knockdown of BRD4 alone did not impact cIAP1 levels, implying that cIAP1 was specifically modulated by the BRD4 degradation products (Fig. 1e). Tracking the kinetics of cIAP1 reduction in MZ1-treated 22Rv1 cells revealed that cIAP1 reduction occurred at 16 hours post-treatment and was accompanied by peak activation of caspase-8, a critical cIAP1-regulated initiator caspase (Fig. 1f)^23^. In the BRD4-L knockdown background, MZ1 failed to reduce cIAP1 and to fully activate both caspase-8 and caspase-3 (Fig. 1f). Other BETd compounds, including VHL-dependent ARV-771^8^ and CRBN-dependent dBET1^4^, dBET6^9^ and BETd-260^26^ (Supplementary Fig. 1), also reduced cIAP1 levels, that were rescued by BRD4-L knockdown (Fig. 1g). These data implicate a common consequence of BRD4-L degradation regardless of E3 co-optation. Taken together, our data indicate that BETd molecules induce caspase activation through the production of specific degradation products that modulate IAP proteins, and not simply by removing BET proteins.

### Proteasomal processing of BRD4-L exposes internal IAP-binding motifs (IBM)

Next, we sought to identify the BRD4-L degradation products that modulate IAP proteins. We hypothesized that proteasomal processing of BRD4-L exposes its internal IBMs that function as bona fide IAP modulators in the context of chemically-induced, constitutive BRD4 degradation. Classical IBMs are defined by their affinity for the BIR3 domains of cIAP1/2 and XIAP, and are capable of inducing cIAP1/2 dimerization and activation through IBM-BIR3 interactions^25, 27^. IBMs match a broad consensus tetrapeptide sequence, (NH2)A-X-P-X, where the first position (P1’) shows a strong preference for N-terminally exposed alanine or serine, and the third position (P3’) prefers proline with some leniency, whereas P2’ and P4’ can tolerate a variety of amino acids^28–30^. To identify BRD4-L peptides with N-terminally exposed alanine and serine residues, we developed a reconstituted ubiquitination and degradation assay using purified components, followed by proteomic analysis (Fig. 2a). While a control experiment without the proteasome identified zero peptides, the complete reaction identified 605 unique peptide sequences matching to BRD4-L, among which 47 and 57 carried N-terminally exposed alanine and serine, respectively (Supplementary Table 1). We narrowed the list to 31 peptides by limiting P3’ to proline or alanine (Supplementary Table 2), and utilized surface plasmon resonance (SPR) to quantify their binding affinities to the cIAP1-BIR3 domain, a critical step for inducing conformational changes in cIAP1 that enables its ubiquitin ligase activity. We confirmed cIAP1 binding by 22 of these 31 peptides (Fig. 2b). Specifically, cIAP1-BIR3 binds to BRD4L-371 (the number denotes P1’ in BRD4-L), BRD4L-1122-1, and BRD4L-1122-2 with single-digit micromolar affinity and with similar kinetics to that of the Smac-WT peptide; whereas, the binding of cIAP1-BIR3 to BRD4L-1308 as well as the rest of candidate peptides was weaker, showing double-to-triple digit micromolar affinity, or in a few cases was not detectable (Fig. 2c, Extended Data Fig. 5–6). Thus, like other reported IBMs, high-affinity BRD4L-derived cIAP1 binders do not strictly follow the classical A-X-P-X consensus^28, 30^. Further, these BRD4L-derived proteolysis products induced cIAP1-mediated E2-Ub discharge *in vitro*, demonstrating their competency for activating autocatalytic cIAP1 degradation (Fig. 2d).

**Fig. 2.**
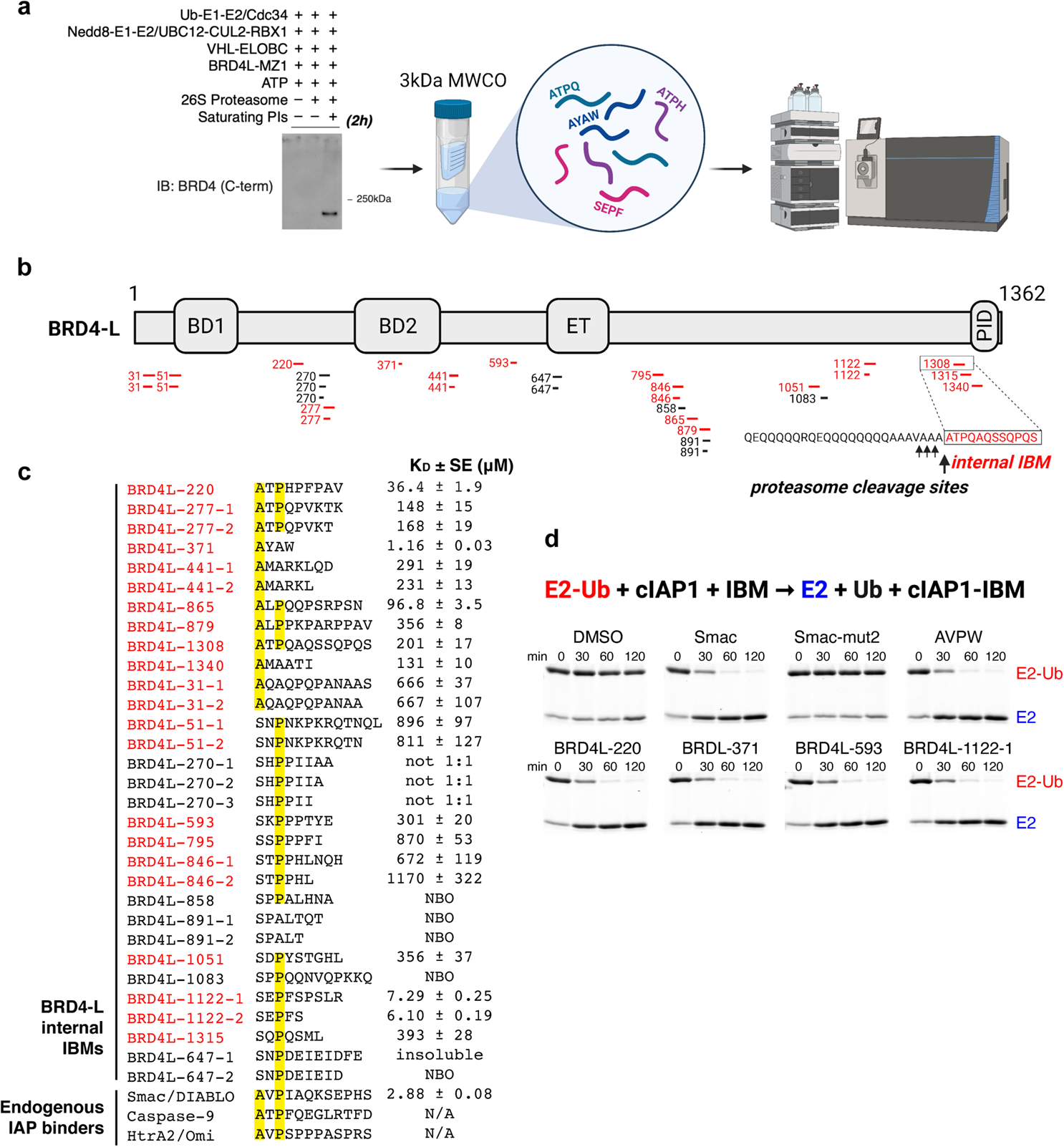
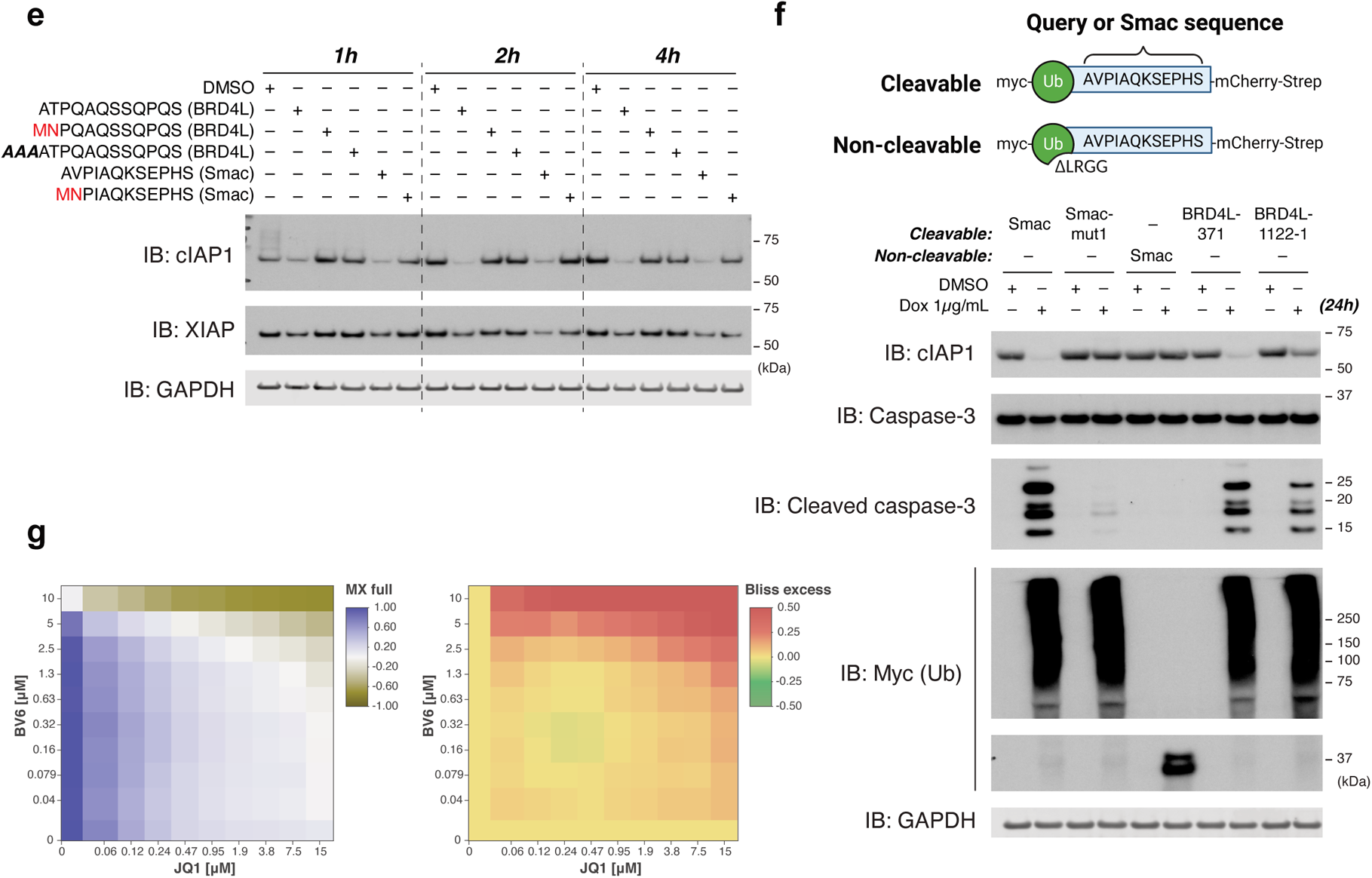
| BRD4-L contains multiple IAP-binding motifs (IBMs) that are liberated by the proteasome, acting as bona fide IAP modulators. **a**, Workflow of *in vitro* ubiquitination and degradation assay followed by isolation of the proteolytic products for mass spectrometry analysis. The saturating proteasome inhibitor cocktail (PIs) is composed of 5 μM bortezomib, 5 μM carfilzomib and 10 μM MG132. BRD4-L degradation efficiency in the absence and in the presence of saturating PIs was examined by immunoblot. **b**, Mapping of the identified BRD4L-IBM candidate sequences to BRD4-L. These sequences follow the criteria of (NH2)A|S-X-P|A-X. Colored in red are confirmed IBMs showing sub-mM to μM binding to cIAP1 (also see Fig. 2c). Arrows denote proteasome cleavage sites. **c**, Sequence alignments of BRD4L-IBM candidates and endogenous IAP-binding proteins. The binding affinities (KD) were determined by surface plasmon resonance; SE denotes the error of the fits reported in the Biacore evaluation software. NBO: No Binding Observed (flat sensorgram, same as the negative control Smac-mut2); not 1:1 indicates data poorly fit by the expected 1:1 binding model; N/A, not assessed (also see Extended Data Fig. 5–6). **d**, IBM-induced cIAP1 activation assay by examining deconjugation of the Ub-conjugated E2 enzyme. *Top,* equation showing ligase-mediated discharge of ubiquitin from the UbcH5c C85S conjugates (E2-Ub), in the presence of the BIR3-RING domain of cIAP1 (abbreviated as cIAP1) and IBM, which is supplied in the form of peptides or DMSO as the control. *Bottom,* the reactants were incubated for the time specified and analyzed by coomassie-stained gels. The sequences of Smac and Smac-mut2 are the same as described in Fig. 2e. **e**, 22Rv1 cells transfected with DMSO or the indicated peptide at 100 μM, followed by incubation in culture media for the time specified. Each lane is an independent transfection experiment. MN denotes two mutations at the N-termini of BRD4L and Smac sequences that block IAP binding. **f**, IBM-induced caspase activation in cells. *Top,* schematics showing the experimental design: dox-inducible IBM expression via Ub(K48R) fusion, which is cleaved by endogenous deubiquitinases; the ΔLRGG mutation in Ub prevents the cleavage. *Bottom,* engineered MDA-MB-231 cells were induced for the expression of the indicated constructs and analyzed by immunoblot. Cleaved Ub was re-utilized by cellular Ub conjugation machinery, resulting in the formation of high molecular weight species, which was absent from cells expressing the non-cleavable construct. Smac-mut1 carries a single A to M mutation at the N-terminus of Smac. **g**, GR values of the 22Rv1 cell line in response to the 5-day combinatorial treatment of the pan-IAP antagonist BV6 and the BETi compound JQ1, calculated based on quadruplicate experiments. Left, MX full represents sigmoidal-fit drug response; GR = 1 no effect, GR = 0 cell stasis, and GR < 0 cell killing. Right, positive Bliss-excess values indicate synergy; zero values indicate additive effects. Unless stated otherwise in Reporting Summary, data represent two to three independent experiments. Source data are provided in Supplementary Information.

To assess whether selective exposure of BRD4-derived neo-amino termini modulates cIAP1 in cells, we compared the transfection of BRD4L-1308, a modest cIAP1 binder (Fig. 2c), with an alternative cleavage product that contains three extra alanines at the N terminus (Fig. 2b; Supplementary Table 1). BRD4L-1308 is preceded by a polyglutamine tract and a seven-residue flanking sequence, the combination of which facilitates proteasomal cleavage within the flanking sequence^31^. BRD4L-1308 induced sustained cIAP1 degradation comparable to the control Smac-WT peptide, whereas the alternative sequence was ineffective at inducing cIAP1 loss, as were the BRD4L-1308 and Smac mutants (Fig. 2e). Consistent with previously published studies, XIAP was less sensitive to IBM-induced degradation than cIAP1 (Fig. 2e)^23^. To further examine the functional consequences of BRD4-L-derived IBM exposure in cells, we engineered Ub(K48R)-IBM fusion for doxycycline-inducible expression in the MDA-MB-231 cell line, a strategy used previously to study N-degron pathways^32^. When expressed, Ub(K48R)-IBM is cleaved by endogenous deubiquitinases, leading to a continuous supply of the IBM peptide, while Ub(K48R)ΔLRGG-IBM prevents deubiquitinase-regulated cleavage. Indeed, exposure to the *in vitro*-validated BRD4L-IBMs (Fig. 2c–d) was sufficient to reduce cIAP1 levels and activate caspases, whereas the exposure to a Smac-mutant sequence was insufficient (Fig. 2f). Finally, JQ1 synergizes with the small-molecule pan-IAP antagonist BV6 in promoting cell death (Fig. 2g), further implicating IAP neutralization as a critical component in BETd cytotoxicity. The combination of *in vitro* reconstitution experiments, biophysical characterization, and functional validation define a precise mechanism to explain why BETd compounds enhance cytotoxicity over BET inhibitors (Fig. 1a–b, Extended Data Fig. 1). This mechanism is defined by an ordered sequence of events: by promoting BRD4-L ubiquitination, BETd compounds target BRD4-L for proteasomal proteolysis (step 1), which exposes IAP-binding sequences (IBMs) within the BRD4-L protein (step 2). When a threshold is reached, IBM peptides promote IAP autoubiquitination and degradation^25^ (step 3), at which point caspases are no longer held in check and their activation promotes apoptotic cell death (step 4).

### Lack of IAP modulation underlies the disconnect between MZ1 resistance and largely functional BRD4 degradation

We next performed cellular resistance studies as an unbiased approach to identify cellular effectors that mediate BETd cytotoxicity. To this end, we derived MZ1-resistant populations and separately derived JQ1-resistant populations as controls from parental 22Rv1 cells that underwent prolonged drug treatments (Fig. 3a). MZ1-resistant cells remained sensitive to JQ1, indicating their dependence on BET for long-term survival (Fig. 3b–c). Additionally, we found no significant cross-resistance between MZ1-resistant and JQ1-resistant clones (Fig. 3c, Extended Data Fig. 7a–b). These results are consistent with our TraCe-seq data, that reveal MZ1- and JQ1-resistant clones are transcriptomically distinct subpopulations (Extended Data Fig. 2).

**Fig. 3.**
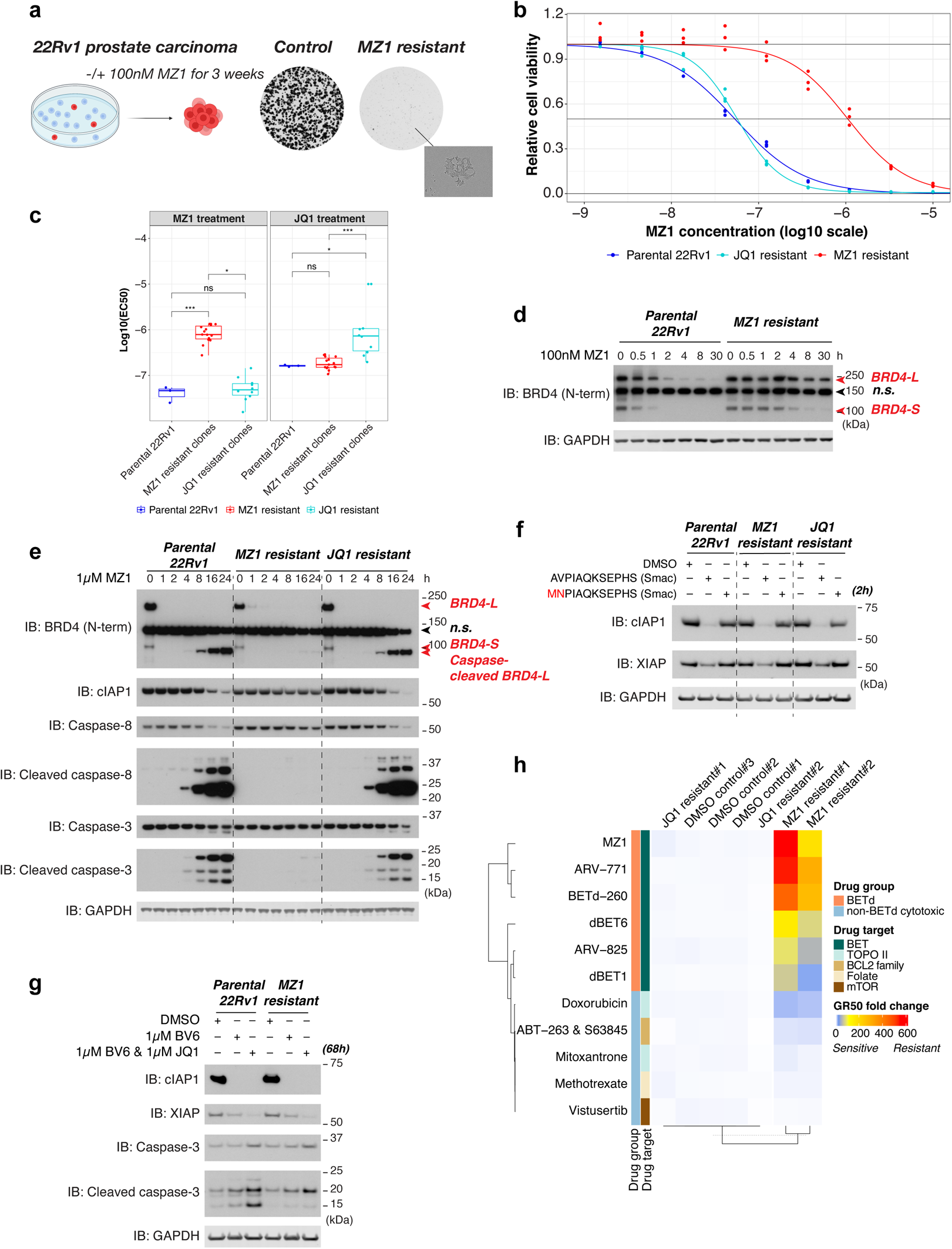
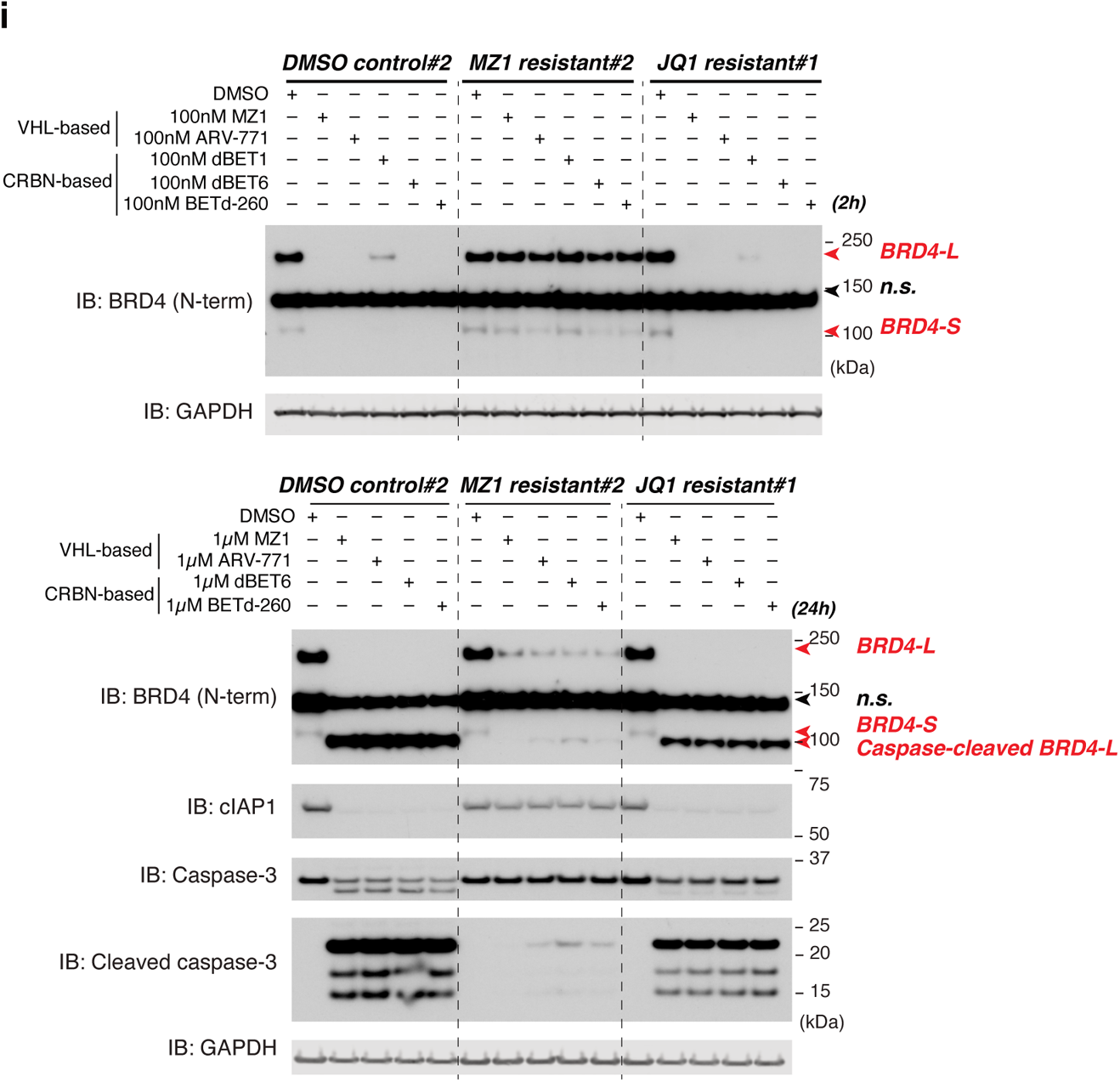
| The lack of IAP modulation contributes to BETd resistance. **a**, Workflow of MZ1 resistance selection in 22Rv1; the expansion of MZ1 resistant clones was monitored by clonogenic survival assay. **b**, MZ1 dose-response curves performed on parental 22Rv1, and bulk-selected MZ1- or JQ1-resistant cells. Data are presented in triplicate experiments. **c**, Log-transformed EC50 for MZ1 and JQ1 treatments in parental 22Rv1, MZ1-, and JQ1-resistant clones, calculated from triplicate experiments. 12 of the MZ1- and 10 of the JQ1-resistant clones were characterized along with the parental line. EC50 is capped at 10 μM for the two JQ1-resistant clones that are non-responsive to JQ1. Box-and-whisker plots show median (solid horizontal line), 25th–75th percentiles (box) and whiskers denote min–max range of the values except for outliers. Adjusted P-values were calculated by unpaired two-sample Wilcoxon test (two-sided) with Bonferroni correction, adjusted P *<0.05, ***<0.001, ns not significant. Also see Extended Data Fig. 7a for experimental attributes. **d,e**, Time course analysis of parental 22Rv1 and bulk-selected resistant cells treated with 100 nM (d) and 1 μM MZ1 (e). **f,g**, Transfection with DMSO or 100 μM of the indicated peptides (f) and treatment of BV6 in combination with JQ1 in parental 22Rv1 and bulk-selected resistant cells (g). **h**, GR50 fold changes for the indicated drug treatments in the independently bulk-selected cells. Data are calculated based on triplicate experiments and normalized to parental 22Rv1. **i**, Immunoblot analysis of the indicated cells in response to 100 nM (i, *top*) and 1 μM BETd compounds (i, *bottom*). Unless stated otherwise in Reporting Summary, data represent at least three independent experiments. Source data are provided in Supplementary Information.

Next, we systematically investigated which step(s) in the BETd-induced cell death response were defective in the MZ1-resistant cells. Despite showing compromised BRD4 degradation when treated with the 100 nM resistance-selection MZ1 concentration, the MZ1-resistant cells showed prompt and near-complete BRD4 degradation when treated with 1 µM MZ1, indicating step 1 remains intact in the setting of MZ1 resistance (Fig. 3d–e). Even with competent BRD4 degradation, cIAP1 levels remained constant and caspases were minimally activated by MZ1 treatment in the resistant cells (Fig. 3e), suggesting a defect in the generation of IBM-containing BRD4 peptides (step 2), or the inability of IAPs to be activated by IBM peptides (step 3), and/or a caspase activation blockade (step 4). IAPs remain functional in MZ1-resistant cells based on their BV6-and IBM-induced degradation (Fig. 3f–g). However, the level of caspase activation induced by BV6, as well as by the JQ1-BV6 combination treatment, was suppressed in these cells (Fig. 3g), indicative of deficits in the generation of IBM peptides (step 2) and the activation of cell death signaling following IAP neutralization (step 4).

Stable MZ1 resistance in drug-free culture conditions suggests that genetic alterations may be partly responsible for MZ1 resistance (Extended Data Fig. 7a). We therefore characterized MZ1-resistant and control clones by whole-exome sequencing. We identified hemizygous or heterozygous mutations in the protein-coding regions of TNFAIP3, RIPK1, the ubiquitin E1 enzyme UBA1 and the proteasomal subunit PSMD11. TNFAIP3 and RIPK1 are critical mediators of the TNFα/NF-κB pathway, the activation of which is the primary mechanism by which pan-IAP antagonists induce apoptosis^23, 24^ (Supplementary Fig. 3–4; Supplementary Table 3; Supplementary Discussion). No genetic alterations were found in any of the BET proteins, IAP proteins, or components of the CRL2^VHL^ ligase. While the combination of UBA1 and PSMD11 alterations may attenuate the rate of BRD4 proteolysis or alter the proteasomal cleavage patterns of BRD4, depletion of either protein alone does not affect MZ1-induced BRD4 degradation (data not shown), highlighting the robustness of the ubiquitin-proteasome system. These data also corroborate our observation that a higher dose of MZ1 enables BRD4 degradation in MZ1-resistant cells (Fig. 3e).

These data predict cross-resistance to other BETd molecules. Indeed, two separate pools of MZ1-resistant cells showed strong and specific cross-resistance to highly potent BETd compounds, and remained sensitive to non-BETd agents (Fig. 3h). Immunoblot analysis confirmed attenuated BET degradation and lack of cIAP1 modulation at low and high doses of BETd compounds, respectively (Fig. 3i). Taken together, these results suggest that MZ1 resistance may arise from combinatorial effects of attenuated target degradation, lower production of IBM peptides, and dysfunctional cell death signaling. Previous reports indicate that the mechanisms of acquired^33, 34^ or engineered^35^ resistance to targeted BRD4 degradation impact ligase machinery; our collective findings emphasize how these differences converge on the reduced capacity to produce IBM peptides that are needed to fully activate caspases.

### A small molecule screen identifies gain-of-function IAP neutralization as a more generalized consequence of degrader compounds

Given the lenient criteria that define IAP-binding peptides^30^, we hypothesized that the IAP-neutralizing mechanism we defined for BET degraders may be more generally applicable to the proteolyzed cellular targets of other TPD-inducing compounds. To systematically investigate this idea, we established a screening workflow to evaluate a list of published E3-interacting compounds that regulate protein homeostasis (Fig. 4a). We knocked in a HiBiT fusion at the endogenous cIAP1 locus in MDA-MB-231 cells and screened two independent clones for compounds that reproducibly decrease the HiBiT-cIAP1 signal. Then, we validated the mechanism of HiBiT-cIAP1 decrease to confirm that knockdown of the reported cellular target of each compound attenuates caspase-3 activity and rescues endogenous cIAP1 loss, by depleting the source of IBM-containing peptides. Among the 46 test compounds evaluated, our top hits were the RNF114 binder Nimbolide^36^, the CDK9 degrader THAL-SNS-032^37^, two pleiotropic degraders that both target Wee1 kinase^38, 39^, and two CDK12-cyclin K-DDB1 glue molecules that target cyclin K for degradation^40, 41^. Importantly, all BET degraders in the screen reduced the HiBiT-cIAP1 signal in a dose-dependent manner (Supplementary Table 4). Knockdown of RNF114, a validated target of the natural product Nimbolide, suppressed Nimbolide-induced endogenous cIAP1 reduction and caspase-3 activation (Fig. 4b). Knockdown of the reported targets of each of the other degrader compounds failed to rescue cIAP1 protein levels and to attenuate caspase-3 activation (Extended Data Fig. 8a–d), perhaps because these compounds also induce proteolysis of other targets that generate IBM-containing peptides, or because cIAP1 levels are decreased via other cellular mechanisms such as transcriptional regulation^37^, leading to caspase-3 activation. The rescue of Nimbolide-induced cIAP1 loss and attenuation of caspase-3 activation by RNF114 siRNA is consistent with reported data, where RNF114 knockdown rescues Nimbolide-induced anti-proliferative effects^36^. Thus RNF114-dependent target degradation, possibly enabled by a Nimbolide-induced molecular glue mechanism, may be responsible for these phenotypes; by contrast, depletion of RNF114 alone by siRNA does not alter cIAP1 levels or caspase-3 activity (Fig. 4b). These data are reminiscent of the cell viability rescue that is achieved by CRBN knockdown in response to treatment with thalidomide analogs^42^, and clinical resistance that manifests in response to CRBN alterations^43^. We therefore evaluated whether the thalidomide analog CC-92480 (mezigdomide) promotes cytotoxicity by inducing neosubstrate proteolysis and thereby generating IBM-containing peptides in a multiple myeloma cell line. Indeed, CC-92480 treatment decreased cIAP1 protein levels and activated caspases, suggesting an IBM-mediated mechanism (Fig. 4c). Importantly, both CC-92480-induced cIAP1 loss and caspase activation were attenuated with CRBN knockdown (Fig. 4c). In sum, the TPD-induced generation of gain-of-function IBM peptides, that neutralize IAPs and thereby activate caspases to promote cell death, is a more generalized feature of E3-binding compounds that induce neosubstrate proteolysis.

**Fig. 4.**
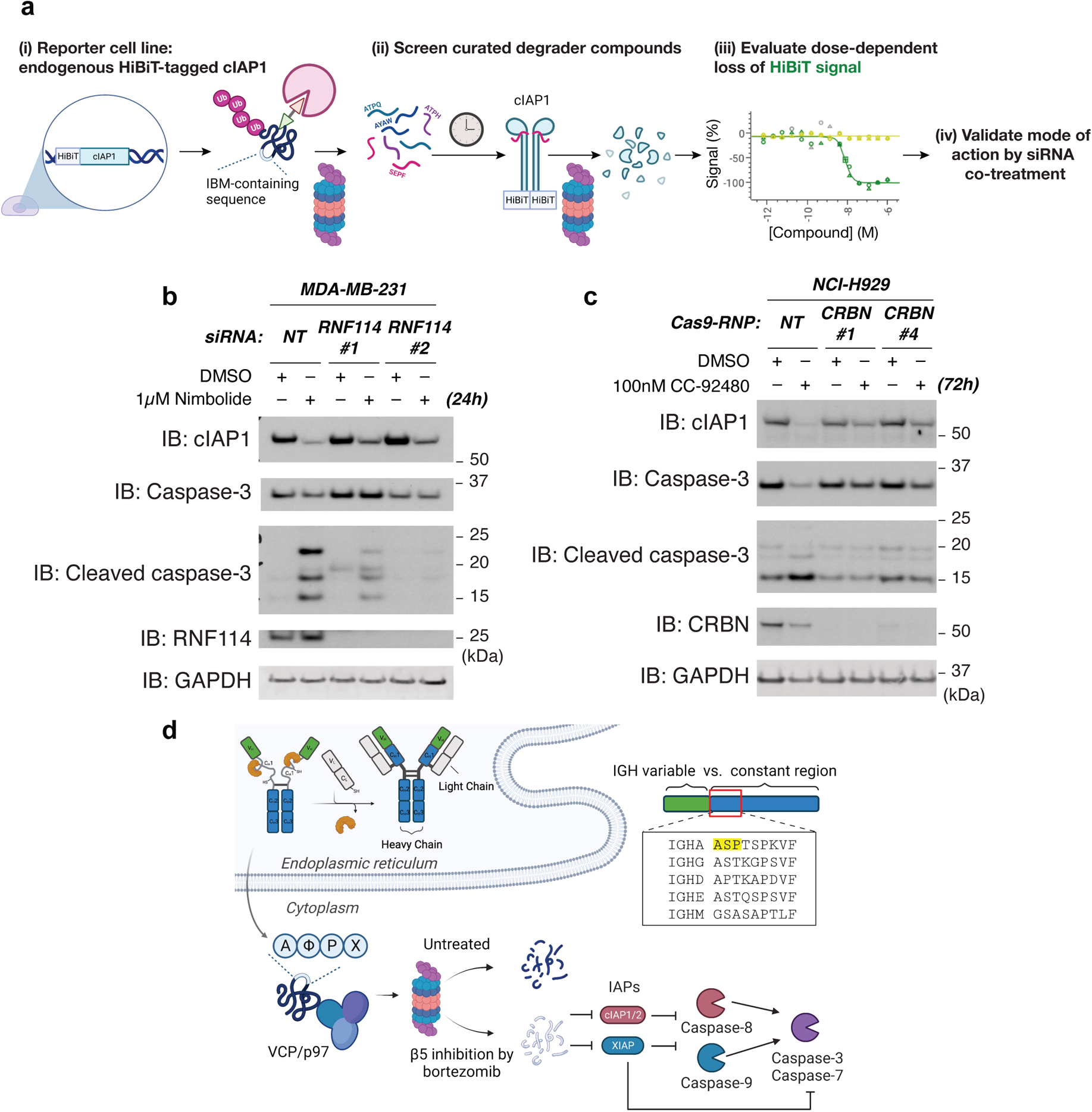
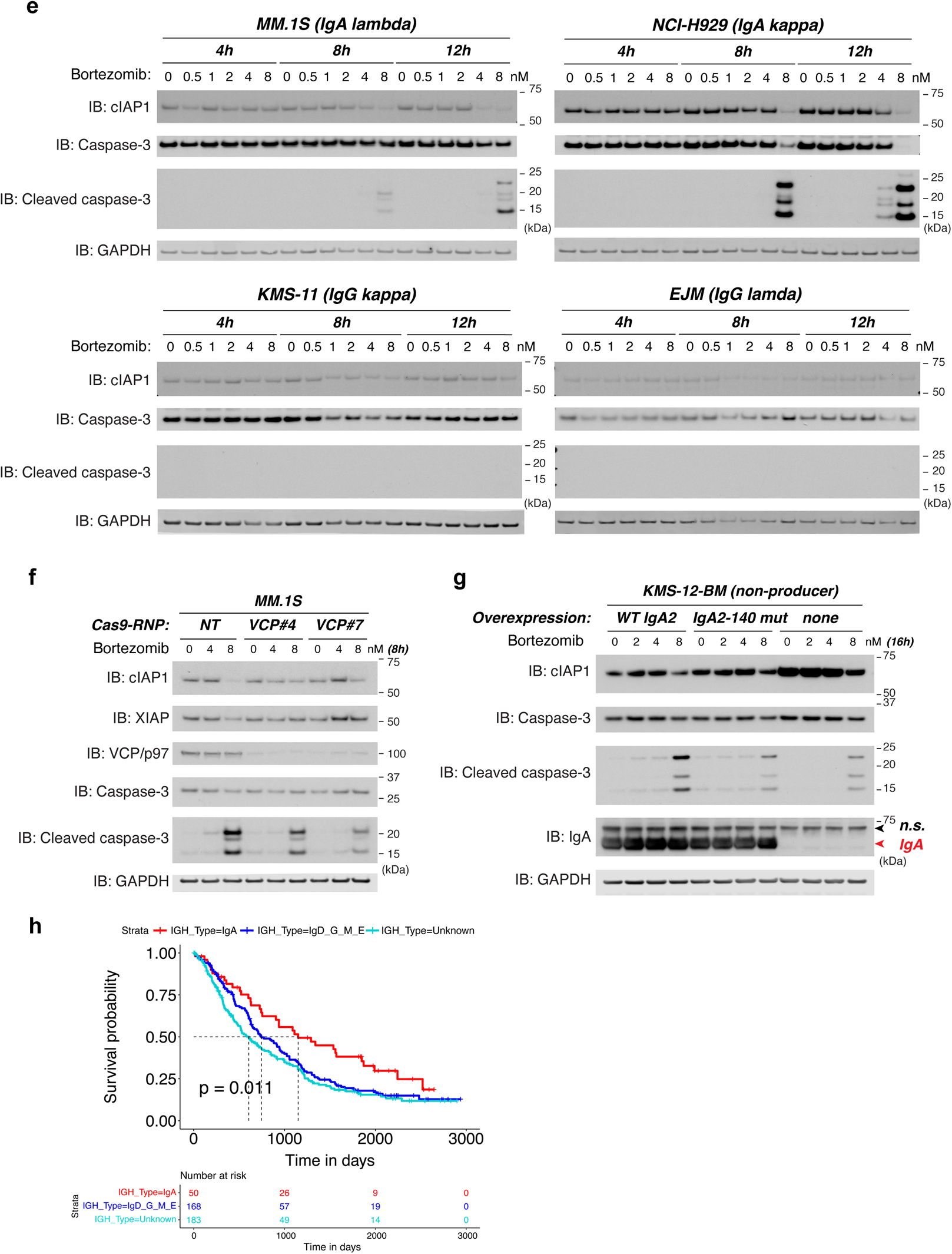
| Modulation of constitutive protein degradation by degraders or limited proteasome inhibition changes proteasome-produced peptide repertoires to generate IBMs, induce cIAP1 loss, and enhance caspase activation. **a**, Screening strategy for degrader molecules that induce cIAP1 loss via IBM-containing degradation products. **b**, MDA-MB-231 cells were transfected with the indicated siRNA for three days and were then treated with the RNF114 recruiter Nimbolide (also see Extended Data Fig. 8a–d). **c**, NCI-H929 cells were transfected with CRISPR-Cas9 ribonucleoprotein (RNP) complex (NT: non-targeting; two separate sgRNAs targeting CRBN), and re-plated two days after transfection to normalize cell numbers, followed by CC-92480 treatments. **d**, Schematics of immunoglobulin (Ig) quality control via ER-associated degradation, in which VCP/p97 extracts and shuttles faulty Ig chains for proteasomal degradation44. IgA harbors a canonical IBM in the constant region that is liberated by a low dose of bortezomib (also see proteomics data in Supplementary Table 5; Extended Data Fig. 9a). **e**, IgA- and IgG-type human multiple myeloma cell lines in response to low doses of bortezomib. All the cell lysates were loaded at 10 μg and developed in the same film cassette for the same period of time for comparison. **f**, Dose responses of bortezomib in control vs. VCP-depleted MM.1S cells. MM.1S was transfected with CRISPR-Cas9 RNP complex (NT: non-targeting; two separate sgRNAs targeting VCP), and re-plated two days after transfection to normalize cell numbers, followed by bortezomib treatments. **g**, KMS-12-BM cells engineered to overexpress wild-type IgA2, IgA2-140 AS-to-MN mutant, or none (parental) were treated with low doses of bortezomib. n.s., non-specific bands. **h**, Kaplan-Meier curves of time to second line therapy of multiple myeloma patients who received bortezomib or bortezomib-thalidomide analog combined therapy without maintenance; stratified by Ig heavy chain (IGH) type: IgA only, other IGH (IgD, IgG, IgM, IgE), and unknown (IGH type was unable to be determined). The pairwise comparison was performed using Log-Rank test and the p-value was corrected for multiple testing (Benjamini-Hochberg method). Unless stated otherwise in Reporting Summary, data represent two to four independent experiments. Source data are provided in Supplementary Information.

### Limited inhibition of the proteasome by bortezomib liberates internal IBMs to activate caspases in IgA-type multiple myeloma

Having demonstrated that heterobifunctional and molecular glue TPD compounds liberate target protein IBMs to activate cell death, we next investigated whether using proteasome inhibitors to directly perturb proteasomal proteases has similar consequences. Bortezomib is a standard-of-care proteasome inhibitor for treatment of multiple myeloma (MM), a malignancy of plasma B cells that relies on endoplasmic reticulum-associated degradation (ERAD) to remove misfolded immunoglobulin chains and to secrete correctly-folded and -assembled antibodies^44^. As such, proteasome inhibition is thought to sensitize MM to cell death due to ERAD perturbation^6^. Bortezomib preferably inhibits the β5 (chymotrypsin-like) and, to a lesser extent, β1 (caspase-like) subunits of 26S/20S proteasome and does not fully inhibit proteasome-dependent proteolysis at clinically relevant doses^45, 46^. The disconnect between incomplete inhibition of proteasomal proteases and potent cytotoxicity by bortezomib remains poorly understood, as does the counterintuitive synergistic combination between bortezomib and standard-of-care thalidomide analogs (e.g. lenalidomide, pomalidomide) that are reported to achieve their therapeutic efficacy by facilitating neosubstrate degradation^1, 2^.

We therefore investigated whether bortezomib produces an alternate peptide repertoire in the context of constitutive antibody degradation in MM cells. If so, differential peptide processing resulting from bortezomib-induced proteasome β5 protease inhibition may expose internal IBMs that neutralize IAPs and activate caspases to promote cell death. To evaluate this idea, we first searched for potential IBMs in the constant region of all five classes of antibody sequences and found that IgA heavy chains (encoded by *IGHA1/2*) uniquely harbor a canonical IBM (Fig. 4d). Further, bortezomib induced loss of cIAP1 and concomitant activation of caspase-3 in IgA-type MM cells at early timepoints and at clinically-relevant concentrations in pre-clinical studies^46^, at which proteasome-mediated degradation was not inhibited *in vitro* (Fig. 4e, Extended Data Fig. 9a–c). Bortezomib-induced cIAP1 loss and caspase activation were not observed in IgG-type cell lines under the same conditions (Fig. 4e). Importantly, both phenotypes in the IgA-type MM.1S cell line were suppressed by prior depletion of VCP/p97, an AAA-ATPase that is critical for extracting and shuttling ERAD substrates for their proteasomal degradation in the cytosol, indicating that bortezomib-induced loss of cIAP1 requires paired VCP/p97 and proteasomal activity (Fig. 4f). Proteomics analysis of *in vitro* degradation products of recombinant IgA by the 26S proteasome confirmed liberation of the predicted internal IBM (IgA2-140), as well as another unpredicted IBM sequence (IgA2-445), in the presence of 10 nM bortezomib but not at a saturating concentration of proteasome inhibitor cocktail (Extended Data Fig. 9a; Supplementary Table 5). Moreover, overexpressing wild-type IgA sensitized the IgA-deficient KMS-12-BM cells to bortezomib-induced cIAP1 loss and caspase activation, supporting the notion that IgA proteolysis promotes bortezomib-induced cell death (Fig. 4g). Expression of the IgA2-140 IBM mutant control showed reduced bortezomib-induced caspase activation, similar to the parental cells (8 nM bortezomib, Fig. 4g). The decrease in cIAP1 levels and elevated background caspase activity by IgA overexpression were attenuated, but not fully abolished with the IgA2-140 IBM mutant (0–4 nM bortezomib, Fig. 4g), likely because the other IgA IBM sequence IgA2-445 revealed by our proteomics data was intact. Thus the extent of cIAP1 reduction and caspase activation induced by bortezomib is proportional to the number of IgA IBMs generated (Fig. 4g; Supplementary Table 5).

These findings prompted us to explore correlations between immunoglobulin heavy chain (IGH) types and survival probabilities of MM patients who received bortezomib therapies. To address this, we analyzed data from newly diagnosed MM patients enrolled in the Multiple Myeloma Research Foundation (MMRF) CoMMpass study. A total of 401 patients who received either bortezomib or bortezomib-thalidomide analog combined therapy as the first line of treatment without maintenance therapy were stratified based on their IGH type. IgA-type MM patients responded to the therapies significantly better than other IGH types or patients of unknown IGH state, with an increase of median time to require second line treatments of 405 and 544 days (1.1 and 1.5 years, p = 0.011), respectively (Fig. 4h; Supplementary Table 6; Extended Data Fig. 10a–c). Our studies reveal that, similar to TPD-inducing compounds, directly perturbing proteostasis in MM cells via partial proteasome protease inhibition also generates gain-of-function IBM peptides. The resultant IAP neutralization and caspase activation lowers the apoptotic threshold to accelerate bortezomib-induced cell death, a mechanism that appears to significantly improve treatment responses of IgA-type MM patients.

## Discussion

Proteolysis-modulating compounds such as thalidomide analogs, proteasome inhibitors, and selective estrogen receptor degraders (SERDs) are standard-of-care treatment regimens for oncology patients. Most therapeutic attention is focused on modulating target proteolysis, whereas the physiological impact of the resulting peptides is often overlooked. Proteasome-generated peptides can achieve gain-of-function status via MHC class I (MHC-I) antigen presentation^47^; our studies now define another gain-of-function role for proteasome-generated peptides in lowering the apoptotic threshold, as governed by IAP proteins^22^. Because IBM peptides have lenient consensus sequences and length requirements^30^, they are likely generated constitutively during homeostatic protein turnover yet have no perceptible effect. Treatment with catalytic TPD compounds, or treatment with proteasome inhibitors to modulate constitutive immunoglobulin proteolysis in multiple myeloma cells, are conditions that generate a robust supply of IBM-containing peptides that exceeds the buffering capacity of IAPs and thus precipitates cell death. Accordingly, it is conceivable that other compounds designed to perturb protein turnover, including p97/VCP, UAE1, and NAE1 inhibitors, may also facilitate cell death by regulating IBM production. Furthermore, IBM generation may be cell-type dependent. Indeed, the thalidomide analog CC-92480 (mezigdomide) promoted CRBN-dependent cIAP1 loss and caspase activation in multiple myeloma (Fig. 4c), but not in MDA-MB-231 breast cancer cells (Supplementary Table 4), presumably due to differentially-expressed neosubstrate proteomes and/or CRBN levels.

Cell death is a therapeutically beneficial outcome of proteolysis-modulating compounds when induced in tumor cells but can lead to dose-limiting toxicity when normal cell types are affected. TPD is expanding beyond oncology to other therapeutic areas^5^, where inducing cytotoxicity within diseased cell types may not be desirable. It will therefore be important to evaluate whether IBM-containing proteolytic products are beneficial, or not, in specific indications, particularly given the context-dependent pro- and anti-inflammatory properties of IAP proteins^48, 49^. To this end, we have outlined a scalable strategy to monitor the proclivity of proteolysis-modulating compounds to generate IBM-containing peptides in any cell type. While much emphasis has been placed on E3 expression analysis to drive tissue-selective target degradation^50^, therapeutic efficacy and toxicity are usually manifest in specific cell types rather than globally in tissues. With the emergence of single-cell analytics, adopting an algorithm that includes cellular sub-type profiling of target, E3, and IAP protein levels combined with target IBM sequence analysis may enable the prediction of therapeutic benefits or toxic liabilities, and thereby inform compound design considerations, to maximize the clinical impact of proteolysis-modulating therapeutics.

## Figure Legends

**Extended Data Fig. 1.**
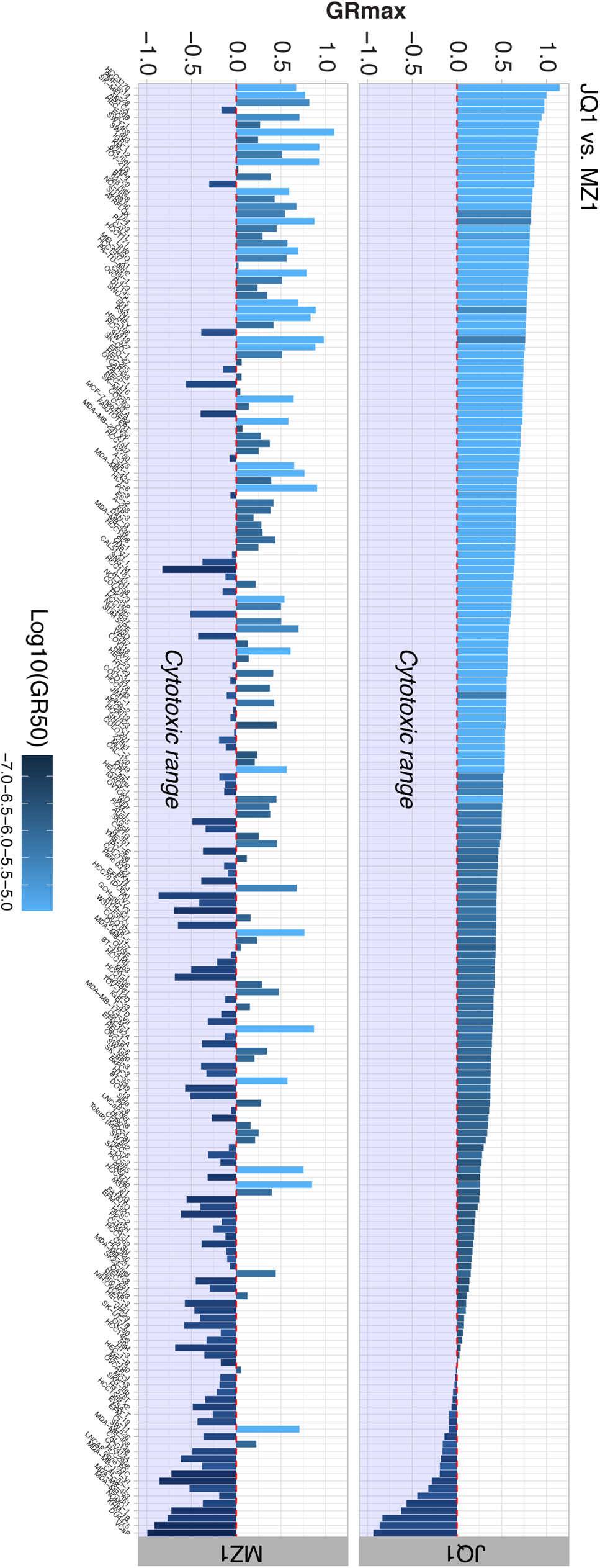
| GR metrics of 192 human cancer cell lines in response to MZ1 and JQ1. As described in Fig. 1a, except that GR50 is capped at 30 µM in cell lines that are not responsive to drug treatments. Source data are provided in Supplementary Information.

**Extended Data Fig. 2.**
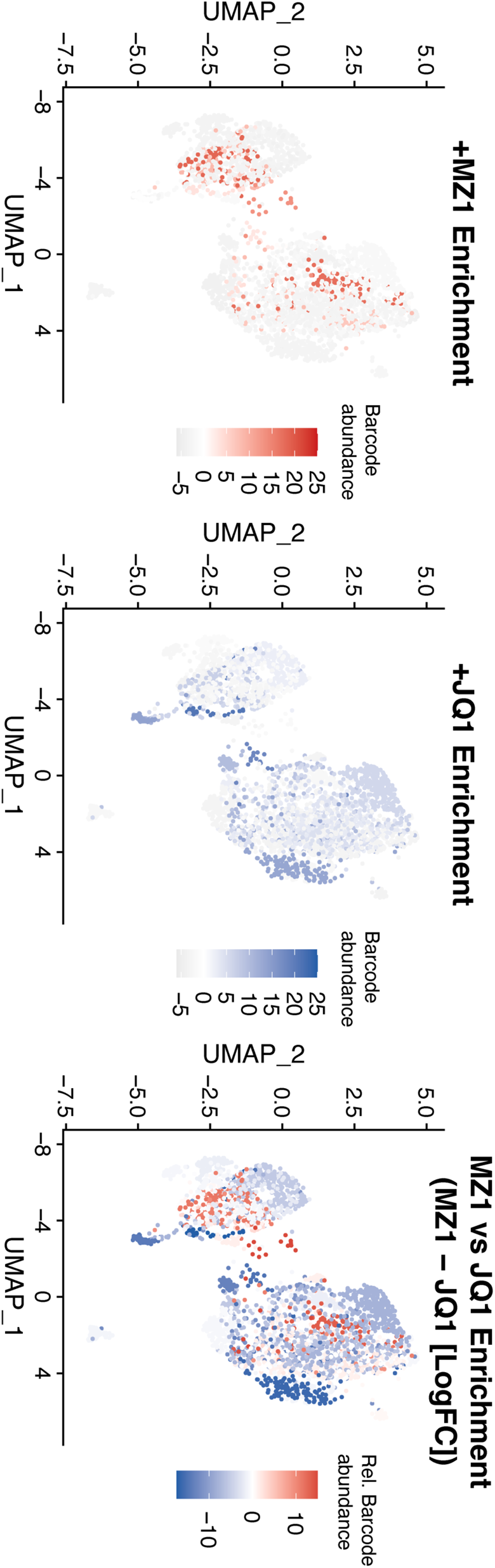
| Single-cell RNA sequencing in combination with clonal tracing reveals little overlap between MZ1 and JQ1 resistant subpopulations. 22Rv1 cell line was engineered to express TraCe-seq barcodes, subjected to single-cell RNA sequencing (baseline transcriptomes), and proceeded with the indicated treatments as previously described^16^. Barcode abundance denotes the enrichment of resistant clonal cells that are presented by their transcriptomic similarities in the UMAP space. Data include four biological replicates of DMSO, three biological replicates of MZ1, and two biological replicates of JQ1 treatments.

**Extended Data Fig. 3.**
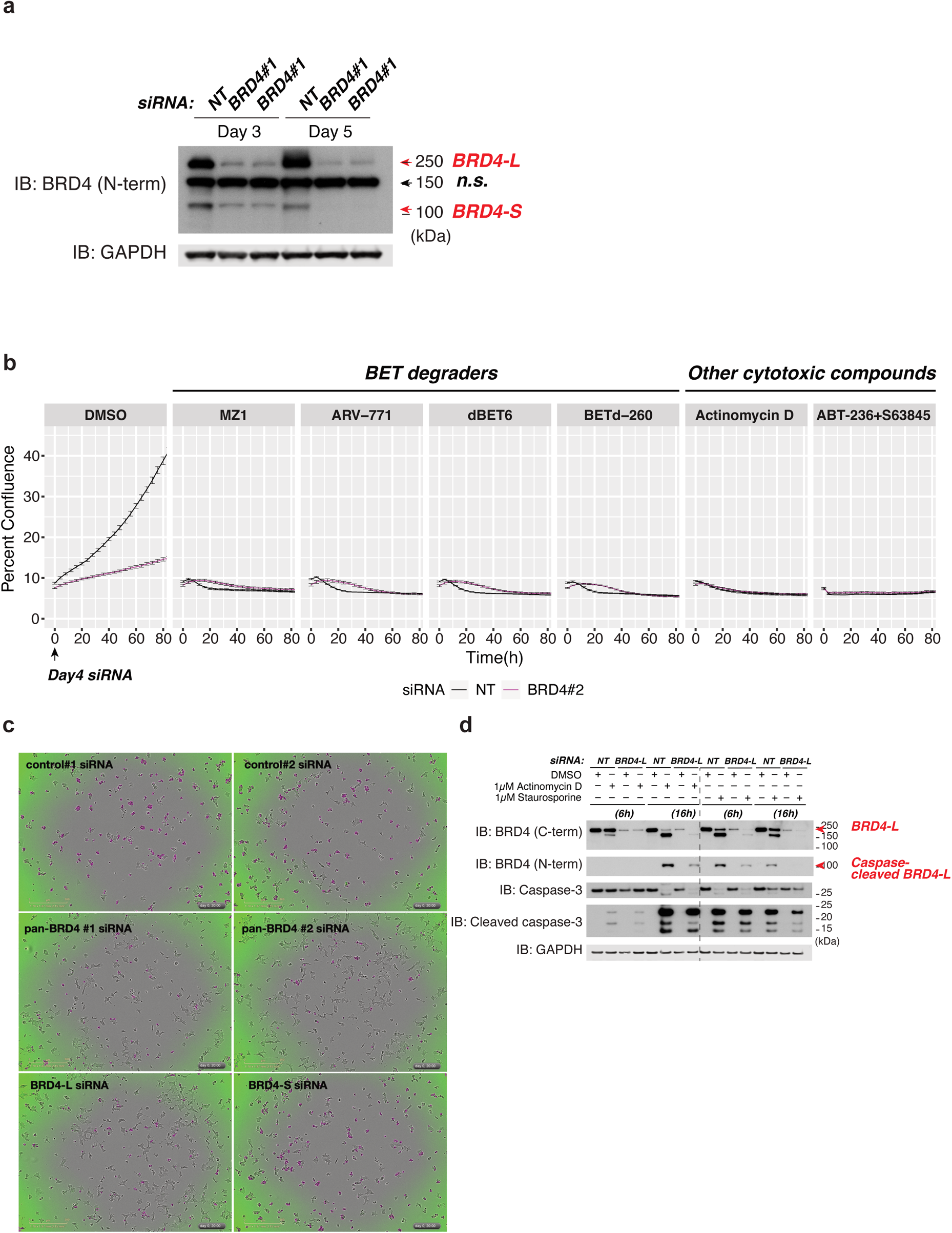
| BRD4 knockdown specifically attenuates the cytotoxicity of BET degraders and not other cytotoxic agents. **a**, Immunoblot to confirm BRD4 depletion. 22Rv1 cells transfected with the indicated siRNA were collected after three days, or re-plated and cultured for additional two days before harvest. **b**, As described in Fig. 1d, time course analysis of percent cell confluence in response to the indicated treatments. Data are presented as mean ± se; n = 3. **c**, As described in Fig. 1d, representative images of siRNA-treated 22Rv1 cells in response to MZ1 treatment at hour 20. Magenta masks denote caspase-3/7 cleavage. **d**, siRNA-transfected 22Rv1 cells were re-plated for the indicated treatments, and cell lysates were analyzed by immunoblot using the indicated antibodies. Unless stated otherwise in Reporting Summary, data represent two to three independent experiments. Source data are provided in Supplementary Information.

**Extended Data Fig. 4.**
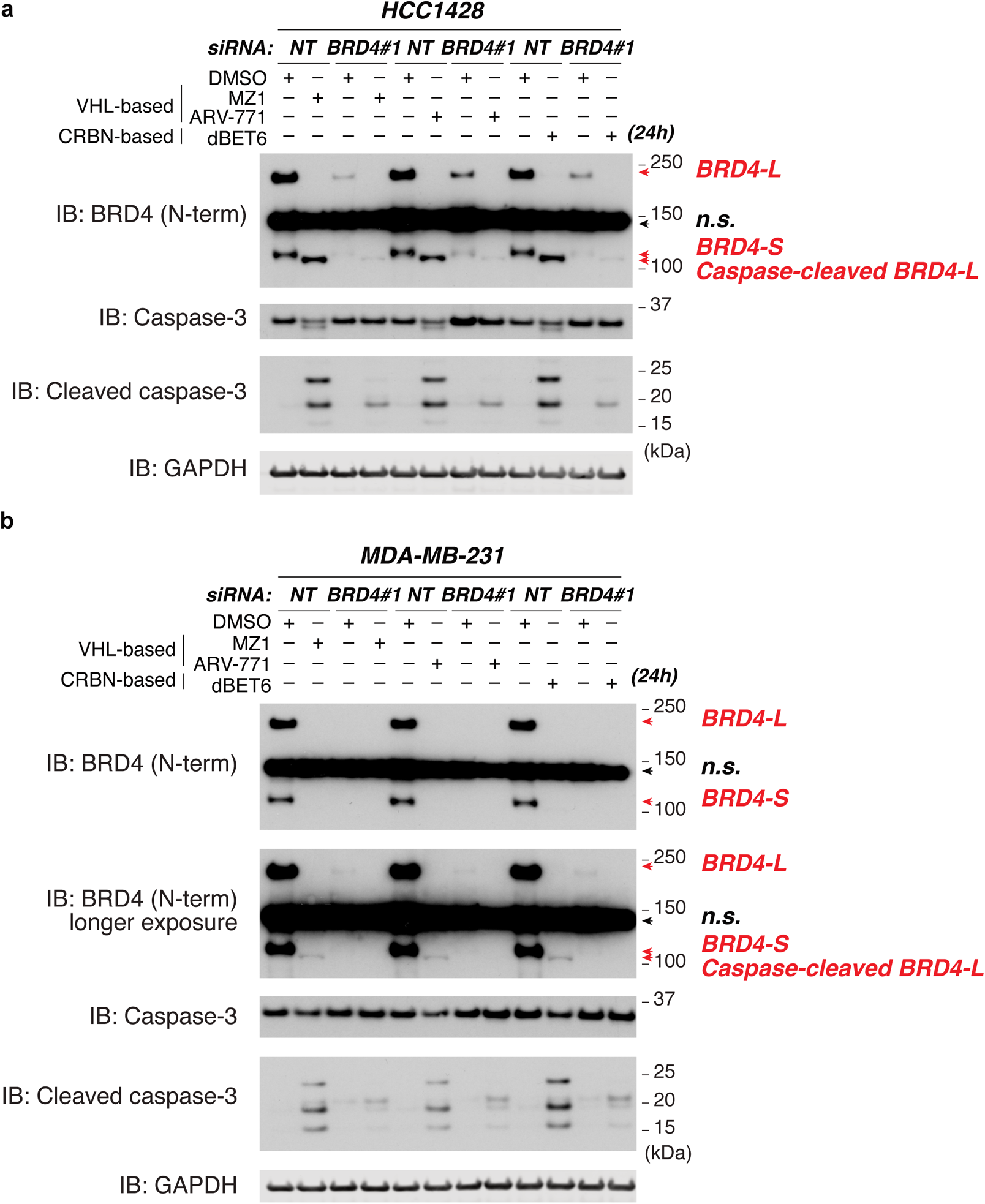
| Rescue of BETd-induced caspase activation in additional cell lines. **a,b**, As described in Fig. 1e, HCC1428 **(a)** and MDA-MB-231 **(b)** cell lines were transfected with siRNA and re-plated for the indicated treatments, and cell lysates were analyzed by immunoblot using the indicated antibodies. Data represent two independent experiments. Source data are provided in Supplementary Information.

**Extended Data Fig. 5.**
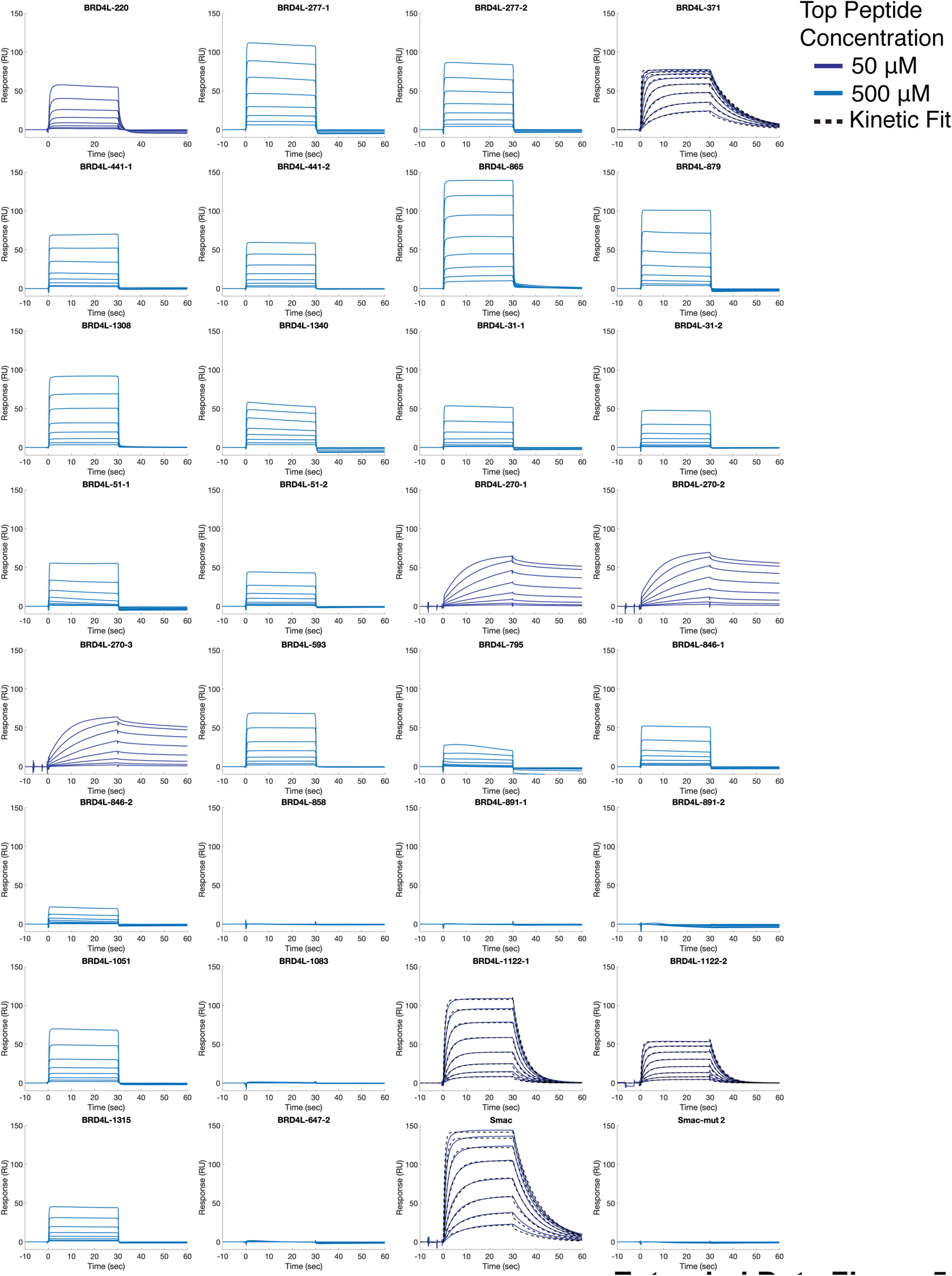
| Validation of BRD4L-IBM candidates by surface plasmon resonance (SPR). SPR sensorgrams of the indicated peptides binding to the cIAP1-BIR3 domain immobilized on the sensor. The derived KD values are reported in Fig. 2c and Extended Data Fig. 6. The dark blue and light blue traces represent two-fold titrations of peptides starting from 50 µM and 500 µM, respectively. The black dashed lines are the kinetic fits from the Biacore S200 Evaluation Software. Data represent two independent experiments. Source data are provided in Supplementary Information.

**Extended Data Fig. 6.**
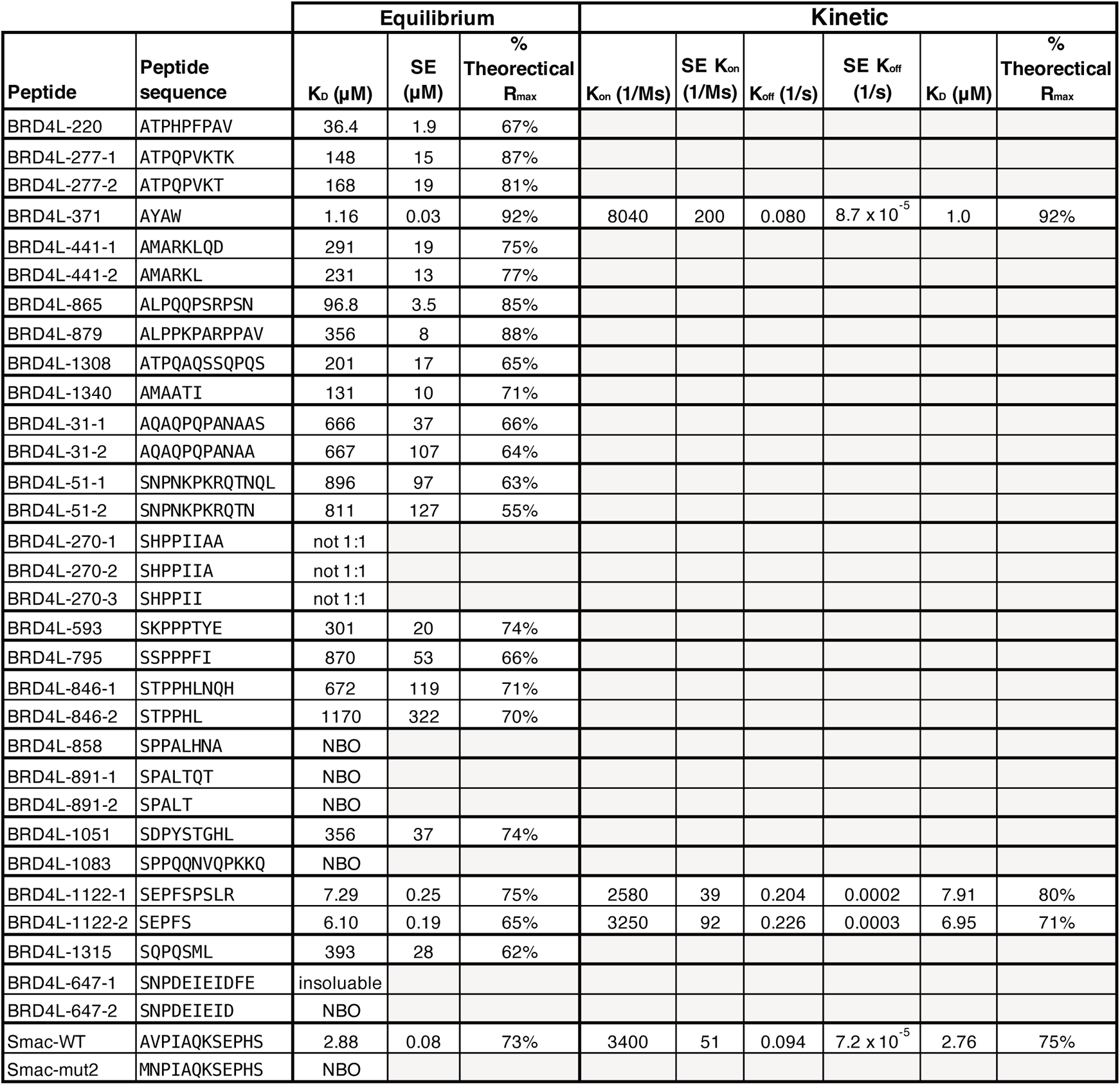
| SPR summary table with kinetic fits. Continued on Extended Data Fig. 5, BRD4L-371, BRD4L-1122-1, BRD4L-1122-2 and Smac-WT displayed good kinetic fits with the derived parameters shown in the table.

**Extended Data Fig. 7.**
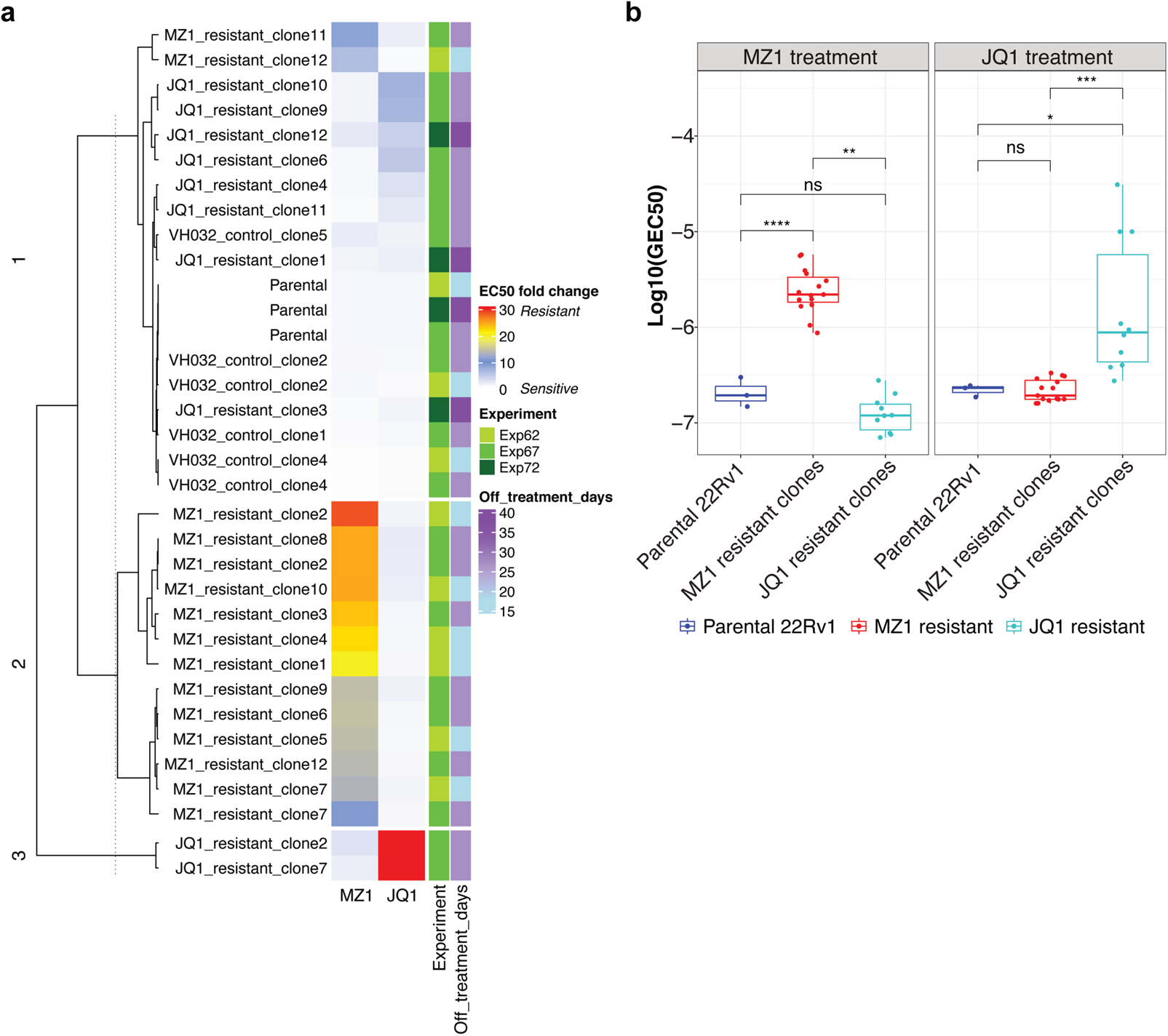
| Drug sensitivity characterization of MZ1-resistant and control clones confirmed no significant cross-resistance. **a**, As described in Fig. 3c, MZ1-resistant and control clones are plotted by their EC50 fold changes in response to MZ1 and JQ1 treatments, relative to the parental 22Rv1; the JQ1-EC50 of JQ1-resistant clones 2 and 7 is capped at 5 µM for the ease of visualization (fold change = 32.3); other experimental attributes including the number of days in drug-free media prior to the treatments and experiment ID are color-coded. **b**, Log-transformed GEC50 (GR-based EC50) is plotted for parental 22Rv1 and the resistant clones. All drug sensitivity metrics are computed based on triplicate experiments. Unpaired two-sample Wilcoxon test (two-sided) with Bonferroni correction, adjusted *P* *<0.05, **<0.01, ***<0.001, ****<0.0001, ns not significant. Source data are provided in Supplementary Information.

**Extended Data Fig. 8.**
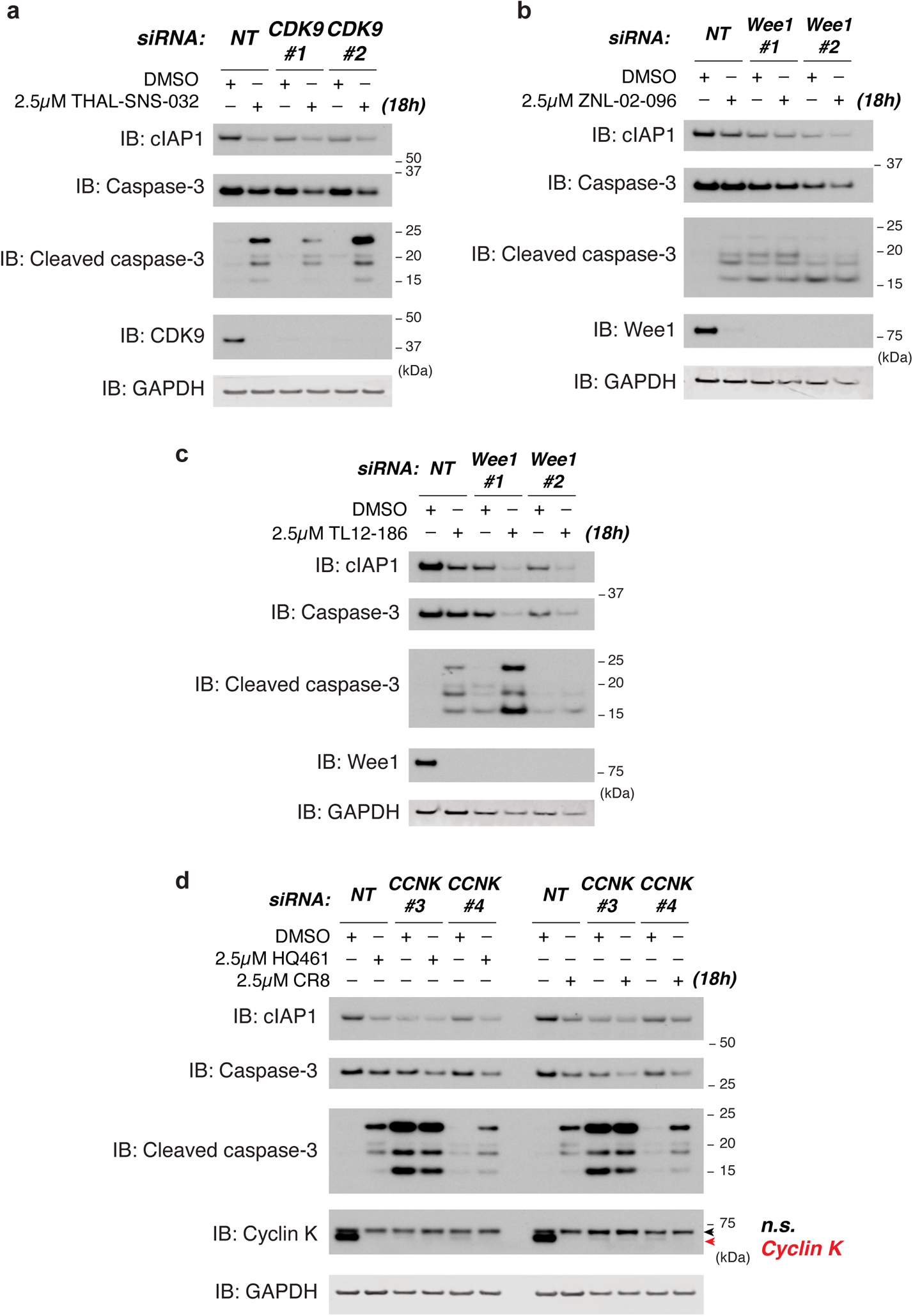
| Validation of degrader hits that reduce cIAP1 levels. MDA-MB-231 cells were transfected with the indicated siRNA for three days and were then treated with the CDK9 degrader THAL-SNS-032 **(a)**, pleiotropic degraders that target Wee1 kinase **(b,c)**, and CDK12-cyclin K-DDB1 glue molecules HQ461 and CR8 that target cyclin K for degradation **(d)**.

**Extended Data Fig. 9.**
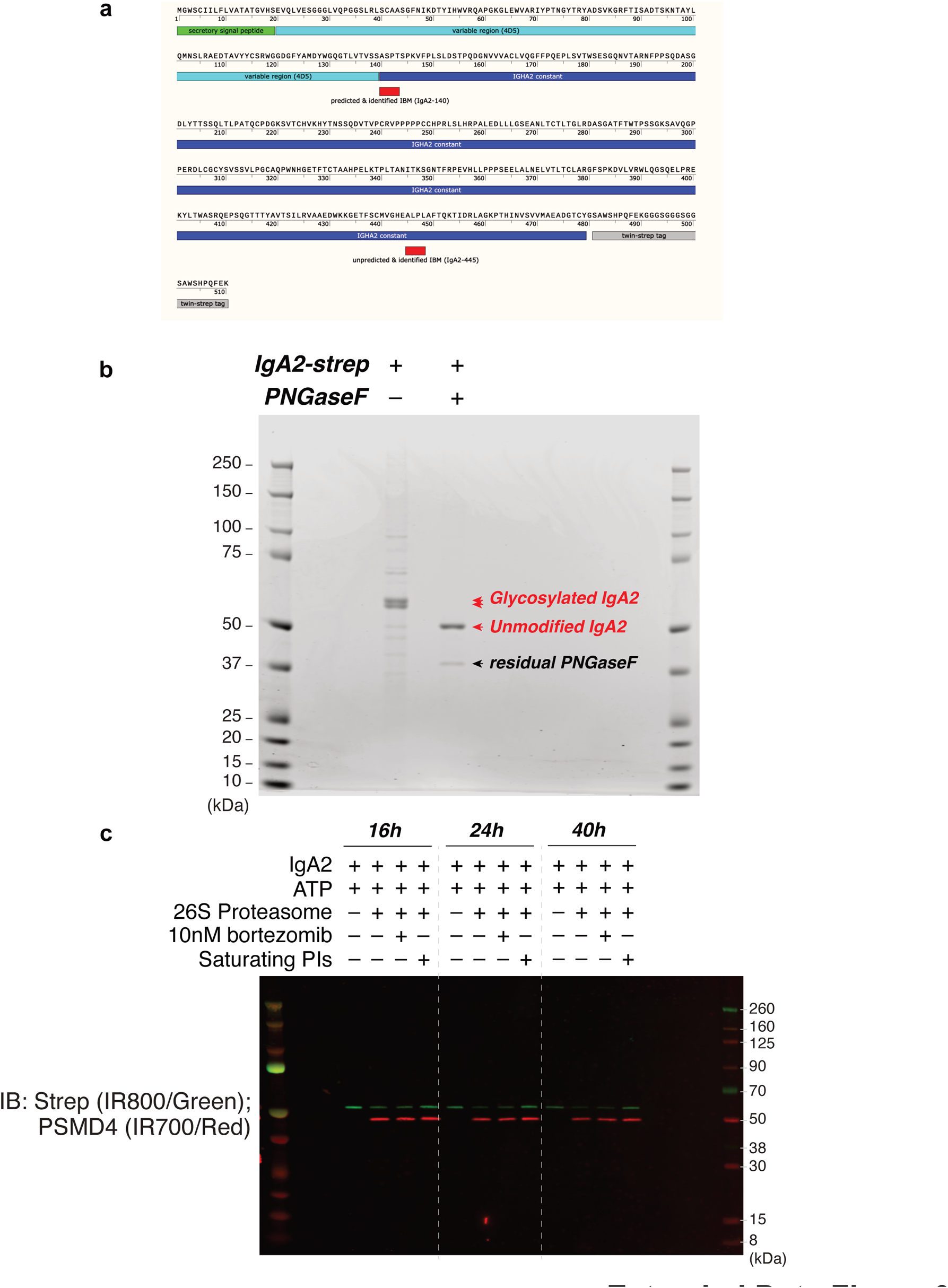
| IgA construct, assessment of IgA purification, de-glycosylation and *in vitro* degradation by purified 26S proteasome. **a**, The design of IgA construct. C-terminally Strep-tagged IGHA2 (IgA2 constant region) is fused with 4D5 (trastuzumab, anti-HER2) heavy-chain variable domain (4D5.VH) as well as a secretory signal peptide at the N terminus. The preceding residue of IGHA1/2 is always serine due to the sequence conservation among the C termini of IGHJ1–6. **b**, Coomassie blue-stained gel image showing Strep-purified IgA2 that is predominantly glycosylated; the purified IgA2 is de-glycosylated by PNGaseF and is subjected to another Strep purification in 3.3 M urea. **c**, *In vitro* degradation of IgA2 with purified 26S proteasome, incubated for the time specified. The saturating proteasome inhibitor cocktail (PIs) is composed of 5 µM bortezomib, 5 µM carfilzomib and 10 µM MG132. Unless stated otherwise in Reporting Summary, data represent at least two independent experiments.

**Extended Data Fig. 10.**
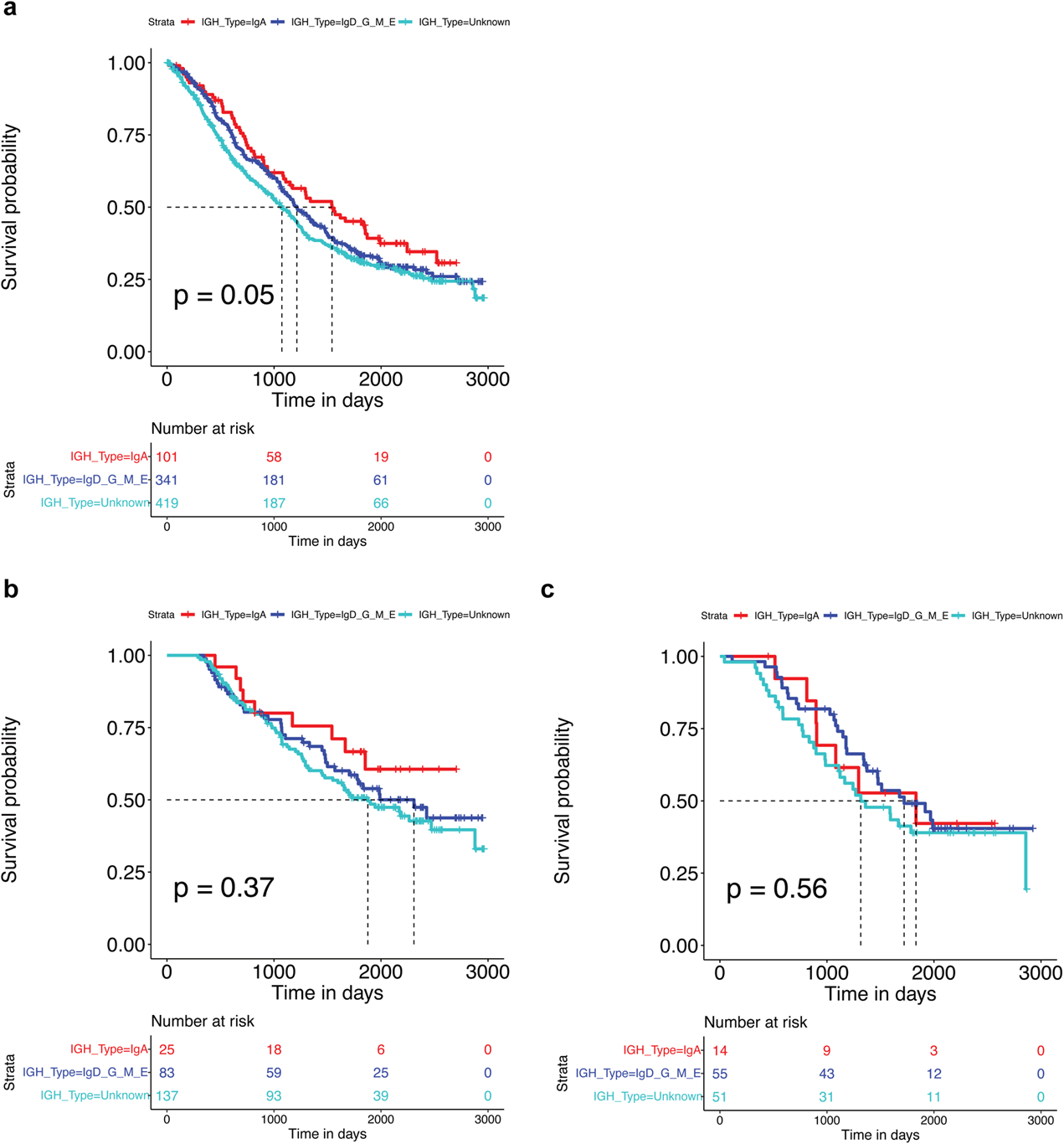
| Correlation between IgH types and time to second line therapy in bortezomib-treated multiple myeloma patients. As described in Fig. 4h, **a**, aggregated data for 861 MM patients with or without concurrent standard (lenalidomide) or non-standard (other agents) maintenance therapy along with bortezomib. **b**, only showing patients who received standard (lenalidomide) maintenance therapy along with bortezomib. **c**, only showing patients who received non-standard maintenance therapy along with bortezomib. Also see Supplementary Table 6.

## Methods

### Cell culture

All cell lines used in this study were obtained from and quality-controlled by Genentech cell line repository. EB-3, MCF7, T47D, HCC1419, LNCaP, 22Rv1, U266B1, LP-1, EFM19, MOLM-16, HCC1428, MV-4-11, EOL-1, VCaP, MM.1S, NCI-H929, KMS-11, EJM, and KMS-12-BM were cultured in RPMI-1640 with 10% heat-inactivated FBS, 2 mM L-glutamine and 1X penicillin/streptomycin. MDAMB415, CAMA1 and MDAMB175II were cultured in DMEM with 10% heat-inactivated FBS, 2 mM L-glutamine and 1X penicillin/streptomycin. MDA.PCa.2b was cultured in DMEM/F12 (1:1) with 20% FBS, 10 ng/mL EGF (354052, Corning), 100 pg/mL Hydrocortisone (354203, Corning), 1X insulin-transferrin-sodium selenite (I1884, Sigma) and 1X penicillin/streptomycin. HEK293 and 293FT were cultured in DMEM with 10% heat-inactivated FBS, 2 mM L-glutamine, 1 mM sodium pyruvate, 1X NEM non-essential amino acids (11140050, Gibco) and 1X penicillin/streptomycin. Expi293F was cultured in Expi293 Expression Medium (A1435101, Gibco) in non-baffled shaker flasks. Cell lines were maintained at a 37°C incubator supplemented with 5% CO2 (8% CO2 for Expi293F).

### Immunoblotting and antibodies

Cells were lysed in 6M urea lysis buffer (also containing 0.1% Triton X100, 100 mM Tris-HCl, pH 7.5, 12.5 mM NaCl, 2.5 mM MgCl2) supplemented with cOmplete Protease Inhibitor Cocktail and PhosSTOP tablets (11836170001 & 4906845001, Roche), sonicated on ice for 5 s at 15 watts, and cleared by centrifugation at 20,000 × g for 10 minutes. Protein concentration of the cleared lysates was determined by Precision Red Advanced Protein Assay Reagent (Cytoskeleton Inc.). Equal amounts of total cell lysates were loaded to 4–12% Bis-Tris or 3–8% Tris-Acetate gels (Invitrogen) and transferred to 0.2 µm PVDF membranes via wet tank or semi-dry method (specified in the supplementary information). After transfer, membranes were blocked with 5% non-fat milk in TBST buffer (with 0.05% Tween20); for anti-Strep detection, membranes were further blocked by Biotin Blocking Buffer (IBA) at 1:1000 in TBST, and incubated with the indicated primary antibodies overnight at 4°C or 2–4 hours at room temperature. All immunoblots were either imaged by chemiluminescent reagents (Pierce ECL or Amersham ECL Prime) and film-based detection or scanned by a LICOR imaging system (Odyssey CLx). Antibodies used for immunoblotting are as following: BRD4 N-term (ab128874, Abcam), BRD4 C-term (A700-004, Bethyl), BRD2 (ab139690, Abcam), BRD3 (A302-368A, Bethyl), Caspase-3 (9662, CST), Cleaved caspase-3 (9664, CST), Caspase-8 (ab32397, Abcam), Cleaved caspase-8 (9746, CST), GAPDH (2118, CST), cIAP1 (AF8181, R&D Systems), XIAP (610717, BD), VCP/p97 (ab109240, Abcam), Strep (SAB2702216, Sigma), PSMD4 (3336, CST), IgA (11449-1-AP, Proteintech), RNF114 (HPA021184, Sigma), CDK9 (2316, CST), Cyclin K (A301-939A, Bethyl), Wee1 (4936, CST), CRBN (HPA045910, Sigma). Secondary antibodies were purchased from LICOR Biosciences and Jackson ImmunoResearch. Uncropped images are provided in Supplementary Figure 5.

### Protein depletion by RNAi and CRISPR

For BET siRNA transfection followed by BET degrader treatments, cells were plated at 2– 2.5E5 cells/mL in 60mm dishes and transfected with 20 nM siRNA using Lipofectamine RNAiMAX (13778075, Invitrogen) after cells became settled on the same day; three days after transfection, cells were trypsinized, counted and re-plated to normalize cell numbers, settled overnight and then treated with compounds. For RNF114, CDK9, Wee1 and Cyclin K siRNA transfection followed by respective degrader treatments, cells were plated and transfected in 12-well plates; three days after transfection, the culture media was replaced with fresh media and then supplemented with compounds. For CRISPR-Cas9, 10 µg (60 pmol) of recombinant Cas9 protein (A36498, Invitrogen) was incubated with 180 pmol of sgRNA in a 20 µL reaction for 20 minutes at room temperature; Cas9-RNP was then mixed with 3E6 of MM.1S or NCI-H929 cells in 100 µL of buffer R and electroporated using Invitrogen Neon system at 2000 V, 20 ms, 1 pulse. After electroporation, the cells were cultured in complete RPMI media supplemented with 20% FBS, and were counted and re-plated for experiments after two days (for VCP/p97). siRNAs were purchased from Thermo Fisher: siBRD4#1 (s23901), siBRD4#2 (s23903), siBRD3#4 (s15545), siBRD3#5 (s15546), siBRD2#5 (s12072), siBRD4-L (GCCAAAUGUCUACACAGUAtt), siBRD4-S (UGGACACGGACUCUUAAUAtt), siNT (4390843), siRNF114#1 (s31751), siRNF114#2 (s31752), siCDK9#1 (s2834), siCDK9#2 (s2836), siCCNK#3 (s16798), siCCNK#4 (s16799). sgRNAs were purchased from Synthego and IDT: sgVCP#4 (CAACAAUUAACCGAUUGGGA, Synthego), sgVCP#7 (GUUUGAGAAUGGCUGUUGAU, Synthego), sgCRBN#1 (AUUCUUCCAUAUCAGCACCU, Synthego), sgCRBN#4 (GCACGAUGACGACAGCUGUC, Synthego), sgNT (GCUUAUGGAUUAGCCAAUC, IDT).

### Chemicals and peptides

Small-molecule inhibitors and degraders were obtained from Genentech compound management except for the following: MZ1 and cis-MZ1 (6154 & 6155, TOCRIS), dBET6 (6945, TOCRIS), BETd-260 and HJB97 (HY-101519 & HY-112429, MedChemExpress), bortezomib and carfilzomib (2204 & 15022, CST), MG132 (474790, Calbiochem). Peptides were synthesized or sourced by the Genentech peptide synthesis group.

### Caspase activity assay with time-lapse imaging

Cells transfected with siRNA were re-plated at 5–7.5E3 cells per well in 96-well plates (3904, Corning) day three post-transfection, settled overnight, and treated with 2 µM Caspase-3/7 Green detection reagent (C10423, Invitrogen) along with the specified compounds. Cell confluence and caspase activity were monitored and analyzed by Incucyte imaging system and the ZOOM software (Sartorius), in which masks for confluence and fluorescent objects were adjusted for individual cell lines of interest. Data were exported and plotted in R.

### Peptide transfection

1E6 of 22Rv1 parental and the resistant cells were re-suspended in 100 µL of buffer R and mixed with 100 µM of the indicated peptides, followed by electroporation at 1530 V, 20 ms, 1 pulse using Invitrogen Neon system. Cells were then cultured in compete RPMI media for the time specified.

### Cell viability assay and drug sensitivity metrics

Seeding density was adjusted based on the doubling times of individual cell lines to obtain non-saturating signal at the endpoint, in 96-well (single agents) or 384-well (for drug combination treatments) formats. Cells were treated with serial dilutions of the indicated compound the next day after plating. For hematological cell lines, the cells were treated for 4 days; for solid tumor cell lines, the cells were treated for 5 days. At the end of the assay, the plates and CellTiter-Glo (CTG) reagents were equilibrated at room temperature, and 50 µL or 25 µL of the reconstituted CTG reagents were added to the 96-well or 384-well plates, respectively. After sufficient cell lysis, the luminescent signal was measured by an EnVision Plate Reader (Model 2104, Perkin Elmer).

For single agent treatments, data were analyzed in R using “GRmetrics” package^51^. For drug combinations, data were processed in R using custom scripts based on GR calculation^52^. Unless otherwise specified, independent groups of samples were analyzed using unpaired two-sample Wilcoxon test (two-sided) with the R package “rstatix”.

### Surface plasmon resonance binding experiments

Reagents were prepared as described previously^27^. The experiments were performed with a Biacore S200 (Cytiva). Biotinylated cIAP BIR3 domain was captured to 2200 RUs on a SA chip (Cytiva) in assay buffer (50 mM HEPES, pH 7.2, 200 mM NaCl, 0.5 mM TCEP, 0.005% (w/v) Tween-20, and 5% DMSO) without DMSO. The surface was then blocked with EZ-Link amine-PEG2-biotin (Thermo Scientific). Binding in assay buffer was collected at 20°C and referenced to FC1. A two-fold dilution series of the peptides (top concentration of 50 or 500 µM, eight concentrations) were injected from low to high concentrations with an association time of 30 s and a dissociation time of 30 s at 60 µL/minute. Data were fit using Biacore S200 evaluation software using a simple 1:1 model.

### Ubiquitin discharge assays

Reagents and assay were prepared as described previously^27^. Briefly, 7 µM of the UbcH5c C85S conjugates (E2-Ub) was combined with 2 µM cIAP-B3R with or without 1 mM peptide in reaction buffer (25 mM Tris, pH 8.0, 20 µM ZnCl2, 2 mM DTT, and 1% DMSO). During the incubation at 30°C, 9 µL of the reaction was stopped with 9 µL NuPAGE LDS Sample Buffer (Invitrogen) at 0, 30, 60, and 120 minutes. 12 µL of each sample was loaded on an SDS-PAGE (Novex WedgeWell 10-20% Tris-Glycine gels, Invitrogen), run for 50 minutes at 200 V, and visualized with InstantBlue Coomassie Protein Stain (Novus Biologicals). Images were taken on the Typhoon 5 (Cytiva) using the IRshort 720BP20 filter.

### TraCe-seq processing and analysis

#### Barcoded cells and treatments

TraCe-seq experiments were conducted similar to previously described^16^. Briefly, 1E7 of 22Rv1 cells were transduced with a TraCe-seq library with 1E5 of barcode complexity at MOI = 0.1. Cells were selected with puromycin and sorted by flow cytometry to enrich for top 50% GFP expressing cells; this is considered TraCe-seq parent barcoded pool. The TraCe-seq parent barcoded pool was cultured for one passage in culture, trypsinized, and 200–300 cells from the barcoded pool were plated in a single well of a 96-well tissue culture plate to establish a TraCe-seq population with ∼100 clones carrying unique barcodes. This starting population was subsequently expanded for 12–14 doublings over 20 days. At this point, cells were seeded in 6-well culture plates at 1E5 cells per well and allowed for attachment overnight. The next day, two wells of cells were trypsinized and dissociated into single cells and subject to single cell RNA-sequencing. The rest of the wells were treated with 1 µM of JQ1 or 200 nM of MZ1 for two months to select for resistant cells. Media was replenished once every week. At the end of the treatment, genomic DNA was extracted from the resistant cells. The barcode regions were amplified by PCR, and barcode abundance was determined by deep sequencing of the amplicon library.

#### Data analysis

TraCe-seq barcode assignment followed as previously described^16^. To identify enrichment/depletion of TraCe-seq barcodes, we applied the DESeq2^53^ method on the raw barcode counts (plus a single pseudocount), and assigned significance of clones based on a minimum log-fold change of three, a *p*-value less than 0.01, a baseMean value of 1000, and at least eight corresponding cells in the scRNA-seq dataset. scRNA-seq data were processed with Cell Ranger 2.1.0 using mkfastq and count commands, and using default filtering. Expression data were processed on the prebuilt human reference GRCh38 and combined using Seurat v3.0.0 following the standard Seurat workflow. The resulting data were filtered to have gene count between 2000 and 6500, and a percent mitochondrial RNA within the bounds of 2 – 10%. The data were log normalized at a scale factor of 10000 and scaled to regress out percent mitochondrial RNA and G2M cell cycle score (Seurat best practices). The first 50 principal component dimensions were utilized for the RunUMAP as well as FindNeighbors and subsequent Louvain clustering at 0.1 resolution. Differential gene expression between sensitive and resistant populations was calculated using the FindMarkers function with a minimum percent cell expression of 0.005. Resulting log-fold changes were input into GSEA analysis using clusterProfiler^54^ with msigdb sets, max geneset size of 1000 and a standard *p*-value cutoff of 0.05.

### Clonogenic assay

Cells were serial diluted from 20,000 to 625 cells per well in 6-well plates, settled overnight, and then replaced with fresh media with or without 100 nM MZ1. Media containing the appropriate compounds was replenished every 72–96 h. After 3 weeks, the cells were washed once with PBS, fixed and stained with 0.5% crystal violet solution (containing 20% methanol) for 20 minutes, washed with water and allowed to dry before scanning using GelCount (Oxford Optronix). For 22Rv1, the optimal seeding density was determined to be 5,000 cells per well; the images were shown in Fig. 3a.

### Degrader screens and HiBiT assay

MDA-MB-231 clones H2 and C10 carrying endogenously tagged HiBiT-cIAP1 were generated by Thermo Fisher and were used for compound screens. Cell lines were cultured in complete RPMI-1640 (supplemented 10% Premium Grade FBS, 1X L-Glutamine, and 1X penicillin/streptomycin). Compound libraries were prepared in 384 well LDV Echo plate as 10 mM, 1 mM, 0.1 mM and 0.01 mM in DMSO, and were dispensed onto 384 well white tissue culture plates (3570, Corning) in a dose response manner (0.1% DMSO final) via Echo. Cells were then plated at 20,000 cells per well in 25 µL media into the pre-dispensed compound-containing plates using 384 ViaFlo robot. Cells were allowed to settle for 30 min at room temperature and then incubated for 18 hours at 37°C in a 5% CO2 humidified incubator. 5 µL CellTiter-Flour (Promega) was added per well to measure cell viability and compounds toxicity. Following a 1-hour incubation at 37°C in a 5% CO2 humidified incubator, fluorescence was measured using an EnVision plate reader (PerkinElmer). For end-point HiBiT lytic detection, 30 µL of reconstituted reagents containing the Nano-Glo HiBiT lytic substrate at 1:50 and the LgBiT protein at 1:100 were added to each assay well; the plates were incubated at room temperature for 10 minutes on an orbital shaker, briefly span on a microplate centrifuge, and measured for luminescence signal on an EnVision Plate Reader (Perkin Elmer).

### 26S proteasome purification

The procedure for generating a tagged-PSMD14 cell line and purification of 26S proteasome using the engineered cell line was adapted from a published study^55^. Briefly, HEK293 cell line stably expressing human PSMD11 carrying a C-terminal HTBH tag was generated by lentiviral transduction and puromycin selection at 2 µg/mL; the HTBH tag consists of two 6xHis tags sandwiching a TEV cleavage site followed by an Avitag peptide for in-culture biotinylation. The stable cell line was expanded using multi-chamber CellSTACK (3313, Corning), trypsinized, washed with cold PBS, flash frozen and stored at -80°C until purification. Cell pellet was quickly thawed and resuspended in lysis buffer containing 50 mM HEPES, pH 7.5, 50 mM NaCl, 10% glycerol, 5 mM MgCl2, 0.5% NP-40, supplemented with fresh 5 mM ATP, 0.5 mM TCEP, 0.1 mM PMSF and 1X cOmplete Protease Inhibitor Cocktail (11836170001, Roche) at 2-bed volumes of the pellet. The cell suspension was homogenized using a Dounce homogenizer (type B pestle) with 15 strokes, and allowed for complete lysis on ice for 15 minutes; the lysates were cleared by centrifugation at 30,000 × g for 15 minutes. Protein concentration was determined using Pierce 660nm Protein Assay Kit (22662). High Capacity NeutrAvidin agarose (29204, Pierce) was equilibrated with the lysis buffer and incubated with the cleared lysates at approximately 1mL packed beads per 200 mg of total proteins at 4°C overnight on an end-to-end rotator. After incubation, the beads were spun at 500 × g, 5 minutes at 4°C; the unbound lysates were removed and the beads were resuspended in the lysis buffer, transferred to a gravity flow column (Bio-Rad), washed with 20-bed volumes of the lysis buffer, and subsequently washed with 10-bed volumes of TEV buffer (50 mM Tris, pH 8.0, 5% glycerol, 0.5 mM TCEP and 2 mM ATP). After washes, the beads were resuspended in 1.5-bed volumes of TEV buffer plus 300 units of TEV protease, and incubated at 4°C overnight with shaking. The eluate was collected by spinning the column, and quality assessed by SDS-PAGE and the overlay of Native PAGE with in-gel fluorogenic chymotrypsin assay. Purified proteasomes were concentrated using Amicon-Ultra 3K MWCO spin columns (UFC500324, Millipore), exchanged to the storage buffer containing 50 mM Tris-HCl, pH 8.0, 10 % Glycerol, 0.5 mM TCEP, 10 mM MgCl2 and 5 mM ATP, concentration determined, aliquoted, flash frozen and stored at -80°C until use.

### Expression, purification and de-glycosylation of IgA

The expression construct for an engineered IgA2 carrying a C-terminal Twin-Strep tag is as described in Extended Data Fig. 9a. Expi293F cells growing at log phase were seeded at 2E6 per mL in HyCell TransFX-H media (Cytiva) in a non-baffled Erlenmeyer flask, and within 1–2 hours after seeding, transfected with 200 µg of the plasmid for a final culture volume of 200 mL using PEIpro transfection reagent (Polyplus). The transfected Expi293F culture was transferred back to 37°C incubator supplemented with 8% CO2 on a shaking platform set for 150 rpm. Customized NS2 feed media was added to the cell culture 18–24 hours post-transfection. 2.5 µM of the p97 inhibitor CB-5083 (HY-12861, MedChemExpress) was added 6 hours prior to harvest, 5 days after transfection. Cells were collected, washed once with cold PBS, flash frozen and stored at -80°C until purification. Cell pellet was quickly thawed and resuspended in 1X HEPES-Urea lysis buffer containing 20 mM HEPES, pH 7.4, 135 mM NaCl, 1 mM EDTA, 1 mM EGTA, 1% NP40, 0.5% sodium deoxycholate, supplemented with fresh 1mM TCEP, 1X cOmplete Protease Inhibitor Cocktail (11836170001, Roche), 10 mM N-Ethylmaleimide, 5 µg/mL PR-619 (SI9619, LifeSensors), 20 µM MG132 (474790, Calbiochem) and 2.5 M urea, at 2-bed volumes of the pellet. The cell suspension was sonicated on ice for 15 s, 3 times, at 30 watts, and cleared twice by centrifugation at 20,000 × g for total of 20 minutes. Total protein concentration of the cell lysates was determined by BCA Protein Assay Kit (23227, Pierce) using diluted lysates in PBS (1:20– 1:60 dilutions). Strep-Tactin XT 4Flow high-capacity beads (IBA) were equilibrated in the HEPES-Urea lysis buffer and incubated with the cleared lysates at 4°C overnight on an end-to-end rotator. The subsequent purification steps were similar to proteasome purification except HEPES wash buffer was used (20 mM HEPES, pH 7.4, 135 mM NaCl, 1 mM EDTA and 0.25% NP40). The recombinant IgA2 was eluted with 2-bed volumes of 1–2X BXT buffer containing 50–100 mM biotin (IBA), buffer exchanged to HEPES-based buffer and to remove biotin using Amicon-Ultra 3K MWCO spin columns, concentration determined, aliquoted, flash frozen and stored at -80°C until use. To remove IgA2 glycans, 40 µL reactions were prepared containing 20 µg of purified IgA2 in 40 mM DTT solution, and heat-denatured at 55°C for 10 minutes; then the reactions were made to a total volume of 80 µL containing 1 equivalent of GlycoBuffer2 and 3–5 µL of Remove-iT PNGase F (P0706, NEB), incubated at 37°C for 2 hours. After incubation, the reactants were spun at 20,000 × g for 10 minutes; the supernatant was transferred to clean tubes; the pellet was resuspended in 20 mM HEPES, pH 7.4, 135 mM NaCl, 1 mM EDTA, 0.5% NP40 with fresh 5 M urea, spun again, and the supernatant was pooled with the previously cleared reactants; the resuspension process was repeated to recover most of the IgA2 (final urea concentration was approximately 3 M). IgA2 was purified again using Strep-Tactin beads, buffer exchanged to HEPES-based buffer, concentration determined, aliquoted, flash frozen and stored at -80°C until use. Purification quality and de-glycosylation efficiency were assessed by SDS-PAGE and coomassie blue staining.

### *In Vitro* ubiquitination and degradation Assay

#### BRD4-L ubiquitination and degradation

Recombinant proteins were expressed and purified in house or purchased from other sources: full-length BRD4-L (RD-21-153, Reaction Biology Corp), Ubiquitin (U-100H, BostonBiochem), UBA1 (E1 enzyme; in house), UBE2R1/CDC34 (E2 enzyme; in house), VHL-ElonginB-ElonginC (substrate receptor-adaptors; in house), CUL2-RBX1 (Cullin-E2 binder, in house), Nedd8 (UL-812, BostonBiochem), NAE1 (Nedd8 E1 enzyme; E-313-025, BostonBiochem), UBC12 (Nedd8 E2 enzyme; E2-656, BostonBiochem). For a total reaction volume of 200 µL, three sub-steps for ubiquitin activation and E2 conjugation (50 µL), CUL2 activation (50 µL), and proteasome mixture (100 µL) were separately prepared. Step 1 comprised 0.1 µM of UBA1, 1 µM of CDC34, 116 µM of Ubiquitin, 2 mM ATP, and 2 mM TCEP in assay buffer (50 mM HEPES-KOH, pH 7.4, 50 mM KCl, 4 mM MgCl2, and 2.5% glycerol), and was incubated at 30°C for 15 minutes. Step 2 comprised 0.3 µM of NAE1, 1 µM of UBC12, 5 µM of Nedd8, 0.05 µM of CUL2-RBX1, 2 mM ATP, and 2 mM TCEP in the same assay buffer, and was incubated at 30°C for 15 minutes, and followed by the addition of 0.05 µM of VHL-ElonginB-ElonginC, 0.1 µM of BRD4-L, and 1 µM of MZ1. Step 3 comprised 0.05 µM proteasome, 2 mM ATP, and 2 mM TCEP in the assay buffer. Step 1 reactants were combined with Step 2, and then Step 3 mixture, which was incubated at 30°C for the time specified.

#### IgA2 degradation

100 µL of 0.2 µM de-glycosylated IgA2 and 100 µL of 0.05 µM proteasome were prepared in separate tubes in the assay buffer supplemented with 2 mM ATP and 2 mM TCEP. The indicated compounds were added to the proteasome preparation if needed. The reactants were combined and incubated at 30°C for the time specified.

#### Peptide discovery (data acquisition)

Peptidome analysis on the *in vitro* reactions by mass spectrometry was performed by MS Bioworks LLC. Briefly, 100% of each sample was filtered by an Amicon-Ultra 3K MWCO spin cartridge. The flow-through was acidified and subjected to SPE on a Waters µHLB C18 plate. The eluates were reconstituted in 0.1% TFA for analysis. Peptides (50% per sample) were analyzed by nano LC/MS/MS using a Waters NanoAcquity system interfaced to a ThermoFisher Fusion Lumos mass spectrometer. Peptides were loaded on a trapping column and eluted over a 75 μm analytical column at 350 nL/minute; both columns were packed with Luna C18 resin (Phenomenex) in BRD4-L experiments and with XSelect CSH C18 resin (Waters) in IgA2 experiments. A 1-hour gradient was employed. The mass spectrometer was operated in data-dependent mode, with MS and MS/MS performed in the Orbitrap at 60,000 FWHM resolution and 15,000 FWHM resolution, respectively. APD was turned on. The instrument was run with a 3-s cycle for MS and MS/MS.

#### Mass spectrometry data processing

For BRD4-L, data were searched using a local copy of Byonic with the following parameters: Enzyme: None; Database: Swissprot Human (concatenated forward and reverse plus common contaminants); Fixed modification: None; Variable modifications: Oxidation (M), Acetyl (Protein N-terminus), Deamidation (NQ); Mass values: Monoisotopic; Peptide Mass Tolerance: 10 ppm; Fragment Mass Tolerance: 20 ppm; Max Missed Cleavages: 2. Byonic mzID files were parsed into the Scaffold software for validation, filtering and to create a non-redundant list per sample. Data were filtered at 1% protein and peptide level FDR and requiring at least two unique peptides per protein.

For IgA2, data were processed through the MaxQuant software v1.6.2.3 (www.maxquant.org) which served several functions: (1) recalibration of MS data; (2) filtering of database search results at the 1% protein and peptide false discovery rate (FDR); (3) calculation of peak areas for detected peptides and proteins. Data were searched using Andromeda with the following parameters: Enzyme: None; Database: Swissprot Human + Custom IgA2 sequence; Fixed modification: None; Variable modifications: Oxidation (M), Acetyl (Protein N-term); Fragment Mass Tolerance: 20ppm. Pertinent MaxQuant settings were: Peptide FDR 0.01; Protein FDR 0.01; Min. peptide Length 7; Min. razor + unique peptides 1; Min. unique peptides 0; Second Peptides: TRUE (which accounts for two peptides being co-transmitted and producing a composite product ion spectrum). The combined folder in MaxQuant was parsed into Scaffold for visualization.

### Plasmid constructs

The Ub(K48R)-IBM and Ub(K48R)ΔLRGG-IBM expression constructs were generated by gene synthesis and cloned to a Tet-on 3G inducible PiggyBac vector using In-Fusion HD enzyme (Clontech). The IgA2 and its IBM mutant constructs were cloned to a PCDNA3.1 vector under the control of a CMV promoter. For stable expression of IgA2 and the mutant constructs in multiple myeloma cell lines, a pLenti6 vector under the control of a CMV promoter was used for cloning.

### DNA transfection

For generating MDA-MB-231 cell lines that express the inducible Ub(K48R)-IBM and Ub(K48R)ΔLRGG-IBM constructs, 4E5 cells were plated in a 6-well plate, settled overnight, and transfected with 1.5 µg of the indicated plasmid DNA and 0.5 µg of the PiggyBac transposase construct using TransIT-X2 (MIR6004, Mirus) according to the manufacturer’s protocol. Three days after transfection, the cells were re-plated in complete RPMI media containing 1 µg/mL of puromycin, which was replaced every 3–4 days until puromycin-treated, mock-transfected cells completely died out. The derived stable cell lines were maintained in 0.2 µg/mL of puromycin, which was removed during experiments.

### Lentiviral production and transduction

For the lentiviral production with pLenti6 constructs, 6E6 of 293FT cells were plated in 100 mm dishes, settled overnight, and transfected with 5 µg of the expression construct, 3.75 µg of the dR8.9 packaging plasmid (pCAG promoter driving HIV-1 Gag and Pol), and 1.25 µg of the pCMV-VSVG envelope plasmid using TransIt-Lenti (MIR6604, Mirus). Three days after transfection, the culture media containing lentiviral particles were collected, spun at 500 × g for 5 minutes, and passed by 0.45 µm syringe filter. The cell lines of interest were plated at 6E5 cells in a 6-well plate, and transduced with lentiviral particles containing 8 µg/mL of polybrene (TR-1003-G, Millipore), which was replaced with fresh media the next day. Three days after transduction, the cells were re-plated in complete RPMI media containing 8 µg/mL of blasticidin, which was replaced every 3–4 days until blasticidin-treated, mock-transfected cells completely died out. The derived stable cell lines were maintained in 1 µg/mL of blasticidin, which was removed during experiments.

### Survival analysis

Kaplan-Meier curves were generated using the R packages “survival” and “survminer”; the *P*-values were derived from the Log-Rank test comparing the groups.

### Data and code availability

All data that support the findings of this study are provided in Supplementary Information or Source Data folder. Custom codes for data analysis are available upon reasonable request.

#### Acknowledgements

We thank Eva Lin for advice on high-throughput drug sensitivity experiments, Micah Steffek for guidance on SPR assays, Christoph Spiess on IgA biochemical properties and engineering, Danielle Kahl for providing Expi293F splits and protein production protocols, Nicholas Clark (University of Cincinnati) for GR method communications, Bartosz Czech for helping with implementing GR method, Shang-Fan Yu and Geoff Del Rosario on exploratory *in vivo* experiments, Samuel Pollock on exploratory experiments with BRD4 peptide discovery, Domagoj Vucic and Tanya Goncharov for sharing their expertise on IAP biology, Karen Gascoigne for helpful discussion on BETd resistance, Aimin Song and the Genentech Peptide Synthesis Group for peptide synthesis support, Yuxin Liang and the Next Generation Sequencing Group for sequencing support, Richard Jones, Christopher Rose and Tommy Cheung for helpful discussion on peptidome analysis.

## Author contributions

S.-H.C. and I.E.W. conceived the project, and I.E.W. supervised the work which was later co-supervised by W.J.F. and X.Y.. S.-H.C. designed, performed and analyzed comparative cis-MZ1/MZ1 drug sensitivity experiments, BETd-siRNA cotreatment studies, and developed working hypotheses with input from I.E.W.. S.P. purified 26S proteasome and designed *in vitro* ubiquitination-degradation assays, and S.-H.C. performed those experiments with input from S.P.. S.-H.C. and W.J.F. discussed and curated BRD4 peptides for follow-up experiments. E.H. designed, performed and analyzed E2-Ub discharge assays and SPR experiments with guidance from E.C.D.; L.R.K. performed a subset of SPR experiments. C.L.G and N.E. designed and performed HiBiT-cIAP1 screens. R.S. designed sgRNA sequences and coordinated with contract research organizations for sourcing the constructs, and S.-H.C. performed cell engineering experiments with input from R.S.. X.Y. designed and performed single-cell clonal tracing experiments. T.S.-W. analyzed TraCe-seq and exome-seq data with input from R.P. and X.Y.. S.-H.C. performed BV6-JQ1 treatments and M.H. analyzed the data. S.-H.C. selected and characterized BETd/i resistant cells, as well as designed, performed and analyzed cross-resistance or the lack of with input from I.E.W., X.Y. and W.J.F.. S.-H.C. characterized MM cell lines’ response to bortezomib, analyzed immunoglobulin sequences, purified IgA2 and performed the degradation experiments. H.H. performed analysis on multiple myeloma patient data and provided guidance on interpreting the data. S.-H.C. wrote the manuscript with input from I.E.W., X.Y., W.J.F., E.H., E.C.D. and R.P.. S.-H.C., I.E.W., X.Y. and W.J.F. edited the manuscript.

## Competing interest declaration

E.H., C.L.G., L.R.K., R.S., T.S.-W., M.H., R.P., E.C.D., H.H., N.E., W.J.F., and X.Y. are current employees of Genentech/Roche. S.-H.C. is a former Genentech postdoctoral research fellow and a current employee of Bristol Myers Squibb. S.P. is a former employee of Genentech/Roche and a current employee of Nurix Therapeutics. I.E.W. is a former employee of Genentech/Roche and a current employee of Lyterian Therapeutics.

## Supplementary Information

Supplementary Tables are provided separately in csv or xlsx format;

Supplementary Discussion with respect to caspase-cleaved BRD4-L and MZ1-resistant clones; Supplementary Figure Legends, Supplementary Figures 1–4, Supplementary Methods, and Supplementary References;

Supplementary Figure 5 containing all uncropped images of immunoblot and coomassie-stained gels in this paper is provided in a separate pdf file.

**Supplementary Table 1 | Mass spectrometry analysis of BRD4-L degradation products.**

**Supplementary Table 2 | Curation of candidate BRD4-L IBM peptides.**

**Supplementary Table 3 | Whole-exome sequencing analysis of resistant and control 22Rv1 clones.**

**Supplementary Table 4 | Screen of curated degrader compounds using HiBiT-cIAP1 reporter cell lines.**

**Supplementary Table 5 | Mass spectrometry analysis of IgA2 degradation products.**

**Supplementary Table 6 | De-identified MM patient data stratified based on IGH type and maintenance therapy status.**

## Supplementary Discussion

### Defining the caspase-cleaved BRD4-L product observed with BETd treatments

In the course of our studies to compare BETi, BETd, and BET siRNA treatments, we noted that BETd treatments were uniquely accompanied by the appearance of a lower molecular weight species that was recognized by anti-N-terminal BRD4 antibodies (Fig. 1c). Because this species co-occurred with caspase activation, we investigated the relationship between BETd-induced caspase activation and appearance of this biomarker. The non-BETd caspase-activating cytotoxic agents actinomycin D and staurosporine also produced the same biomarker, which was suppressed by co-treatment with the pan-caspase inhibitor Z-VAD-FMK (Supplementary Fig. 2a). Neither BET inhibition nor RNAi-induced BET depletion led to BRD4-L cleavage, consistent with the absence of caspase activation (Fig. 1c). These findings suggest that the observed biomarker is a BRD4-L cleavage product resulting from caspase activation.

In order to verify that the biomarker is indeed derived from BRD4, we monitored the cleavage product and caspase activity in BRD4-depleted cells upon treatment with either BETd or non-BETd cytotoxic agents. These experiments yielded an unexpected finding that the level of caspase activation by BETd was reduced in BRD4-depleted cells, suggesting that the degradation products could be responsible for BETd-induced caspase activation (Fig. 1d). BRD4-L siRNA also reduced the appearance of the BRD4-L cleavage product in both BETd- and non-BETd-treated cells, confirming that the biomarker is derived from BRD4-L and is downstream of caspase activation (Supplementary Fig. 2b). However, only BETd-induced caspase activation is suppressed by prior BRD4 depletion (Extended Data Fig. 3b, 3d). These findings prompted us to investigate the mechanism by which BETd requires the presence of BRD4 to fully activate caspases.

### MZ1 resistance mechanisms in 22Rv1 cells based on genetic alterations

To uncover potential molecular determinants of MZ1 resistance, we performed whole-exome sequencing on three of the MZ1-resistant, three of the JQ1-resistant, as well as the parental 22Rv1, which served as a reference for computing allele frequencies (AF) to inform zygosity.

All three sequenced MZ1-resistant clones simultaneously carried a hemizygous V12M mutation in the X-linked ubiquitin E1 enzyme UBA1, a heterozygous A5V mutation in the proteasomal subunit PSMD11, a heterozygous truncating variant of A20 (encoded by *TNFAIP3*) that lacks the zinc finger domains ZnF4–ZnF7 (i.e. 1–535), and two out of the three clones also carried a heterozygous T38I mutation in the kinase domain of RIPK1. None of these mutations were present in the JQ1-resistant clones (Supplementary Fig. 3). We postulated that the combination of UBA1 and PSMD11 mutations may explain the observed attenuation of MZ1-induced BRD4 degradation in the resistant cells. However, these genetic alterations did not completely abolish global ubiquitin-proteasome system (UPS) function, as demonstrated by near-complete BRD4 degradation in the MZ1-resistant cells when treated with 10-fold higher concentration of MZ1 (Fig. 3d–e, 3i). These results indicate the robustness of the UPS and also imply that the quantities and identities of degradation products could be more important in constitutive target degradation induced by degrader molecules. The latter is supported by the lack of both cIAP1 modulation and caspase activation in these resistant cells despite competent BRD4 degradation (Fig. 3d–e, 3i). Direct evaluation of degrader-induced IBM peptide production in cells is complicated by co-degradation of IBM peptides with cIAP1^56^. Therefore we turn to assessing cIAP1 function and signaling in the resistant cells by treating them with IAP antagonists (Fig. 3f– g). The results indicate that cIAP1 remains functional and yet caspases are minimally activated despite cIAP1 neutralization (Fig. 3f–g). These observations are consistent with our whole-exome sequencing data, in which the MZ1-resistant clones carry wild-type IAP proteins, mutant A20, and mutant RIPK1 (Supplementary Fig. 3). Both A20 and RIPK1 are critical mediators of the TNFα/NF-κB pathway. Specifically, IBM-induced cIAP1/2 degradation stabilizes NF-κB-inducing kinase (NIK), which then induces activation of the non-canonical NF-κB pathway that drives autocrine production of TNFα, which subsequently stimulates TNF receptor-1 (TNF-R1) signaling that, in the absence of cIAP1/2, leads to the formation of RIPK1-dependent apoptosis-inducing complex IIb, or the formation of RIPK1-and-RIPK3-dependent necroptosis-inducing complex when caspase-8 is inhibited^57, 58^. Regulation of RIPK1 kinase activity by phosphorylation in *trans* and *cis* constitutes an important checkpoint for the commitment of RIPK1-dependent cell death, and T38 is a reported phosphorylation site in large-scale proteomic studies^58^. We therefore speculate that the T38I mutation might serve as a protective brake against RIPK1-dependent cell death. The ZnF4 and ZnF7 motifs of A20 bind to K63-linked and linear polyubiquitin chains, respectively^59^. Thus the truncation mutation of A20 lacking both regions cannot be recruited to TNF-R1, leading to enhanced K63-linked polyubiquitination of TNFR1 complex proteins, which then facilitates the assembly of the pro-survival instead of the pro-death NF-κB signaling complexes in the presence of sufficient linear ubiquitination^59, 60^. In sum, our experimental findings, together with established knowledge of TNFα/NF-κB signaling, lead us to propose a model to explain MZ1 resistance in 22Rv1 cells (Supplementary Fig. 4). Although we cannot exclude the involvement of non-genetic factors in conferring MZ1 resistance, the stable maintenance of resistance in MZ1-free culture media for a minimum of 18–38 days, suggests that the identified genetic aberrations play a significant role in MZ1 resistance (Extended Data Fig. 7a).

**Supplementary Fig. 1.**
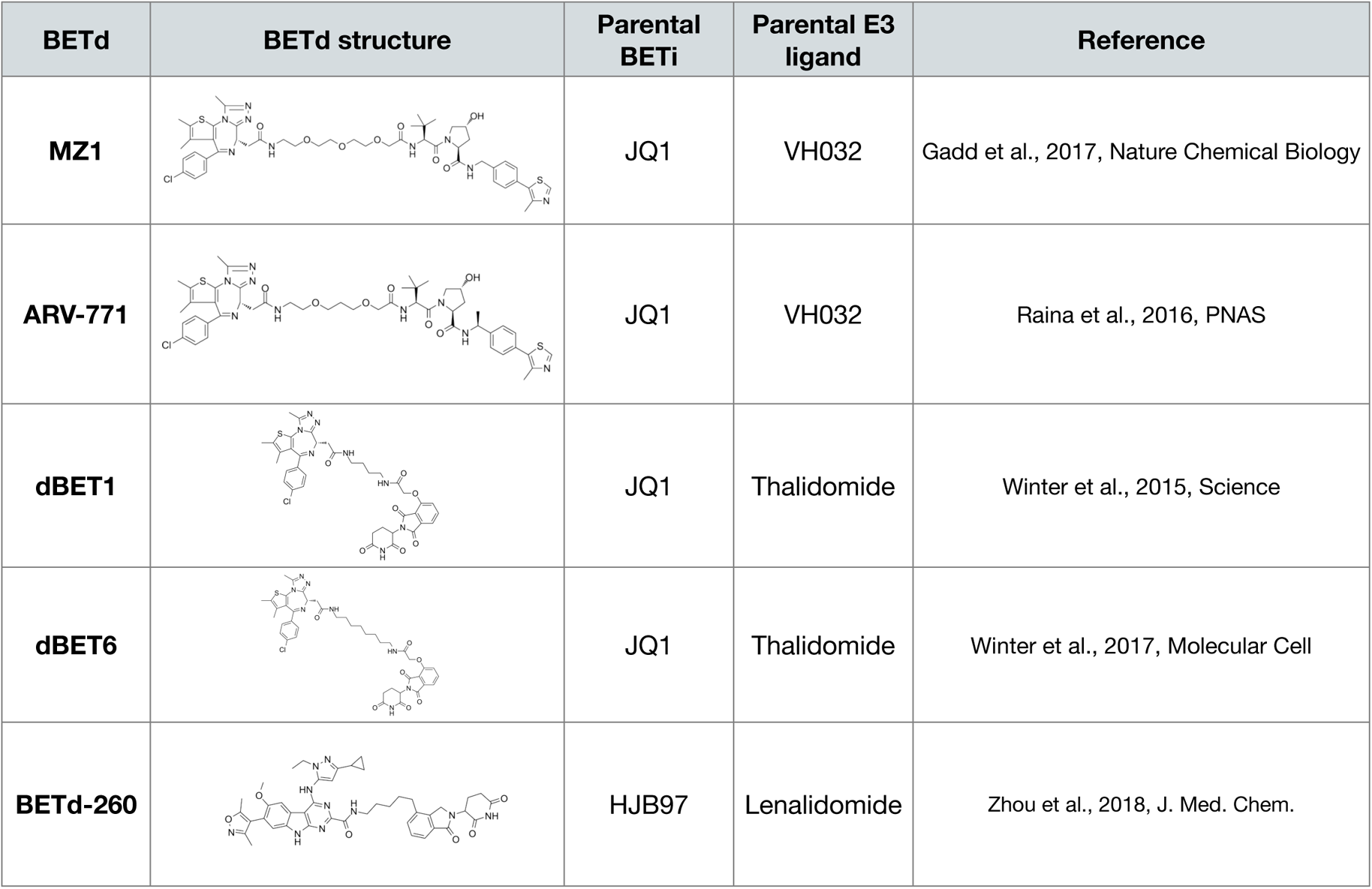
| Chemical structures of BETd and the corresponding parental ligands.

**Supplementary Fig. 2.**
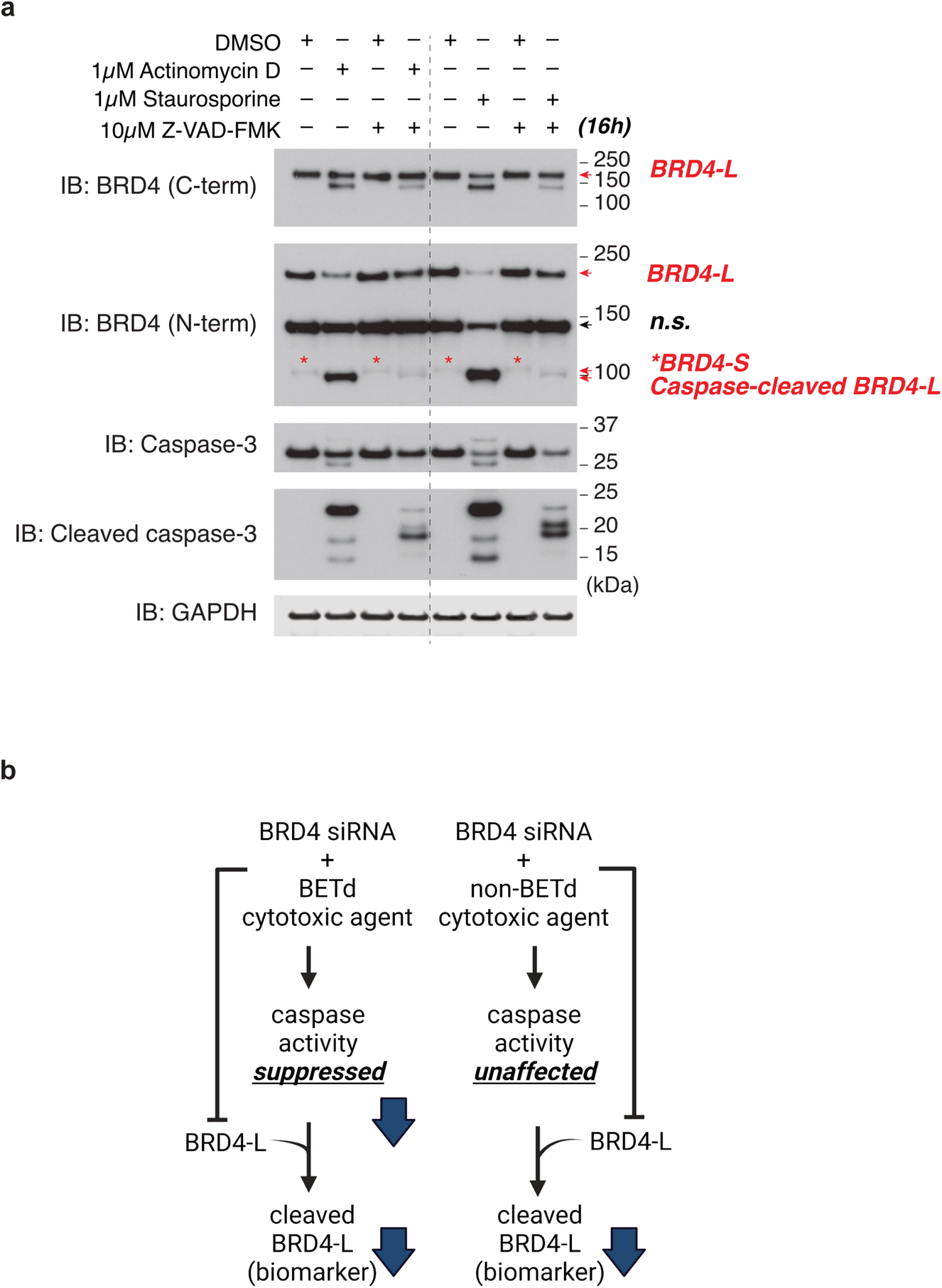
| Cytotoxic agents induce caspase-dependent cleavage of BRD4-L. **a**, 22Rv1 cells treated with DMSO or the indicated cytotoxic agents in the presence or in the absence of the pan-caspase inhibitor Z-VAD-FMK. Red asterisks are centered over BRD4-S, a band distinct from the caspase-cleaved BRD4-L (slightly lower band). Data represent two independent experiments. Source data are provided in Supplementary Information. **b**, Experimental summary of changes in the BRD4-L cleavage product (biomarker for caspase activation) by treatments with BETd or non-BETd cytotoxic agents. BRD4-L siRNA reduces the biomarker, confirming that it is derived from BRD4-L. While BRD4 siRNA does not affect caspase activation by non-BETd cytotoxic agents, it suppresses BETd-induced caspase activation (also see Extended Fig. 3d).

**Supplementary Fig. 3.**
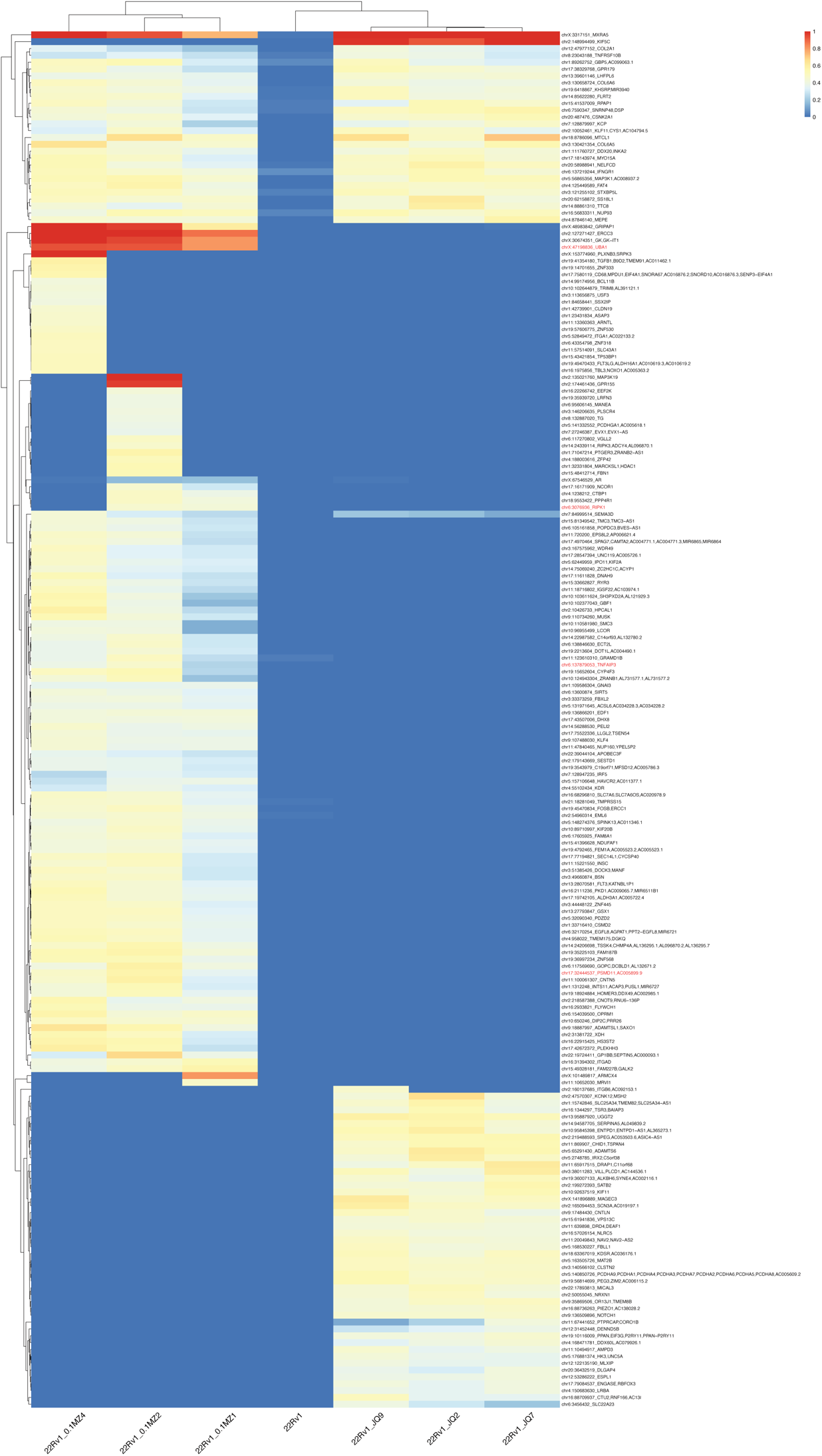
| Protein-coding gene variants identified in MZ1- and JQ1-resistant clones. Heatmap of average allele frequencies (AF) using three different variant calling methods. AF value of 1 denotes homozygous or hemizygous (22Rv1 cell line only carries one X chromosome) mutation, and AF value of 0.5 denotes heterozygous mutation. Also see Supplementary Table 3.

**Supplementary Fig. 4.**
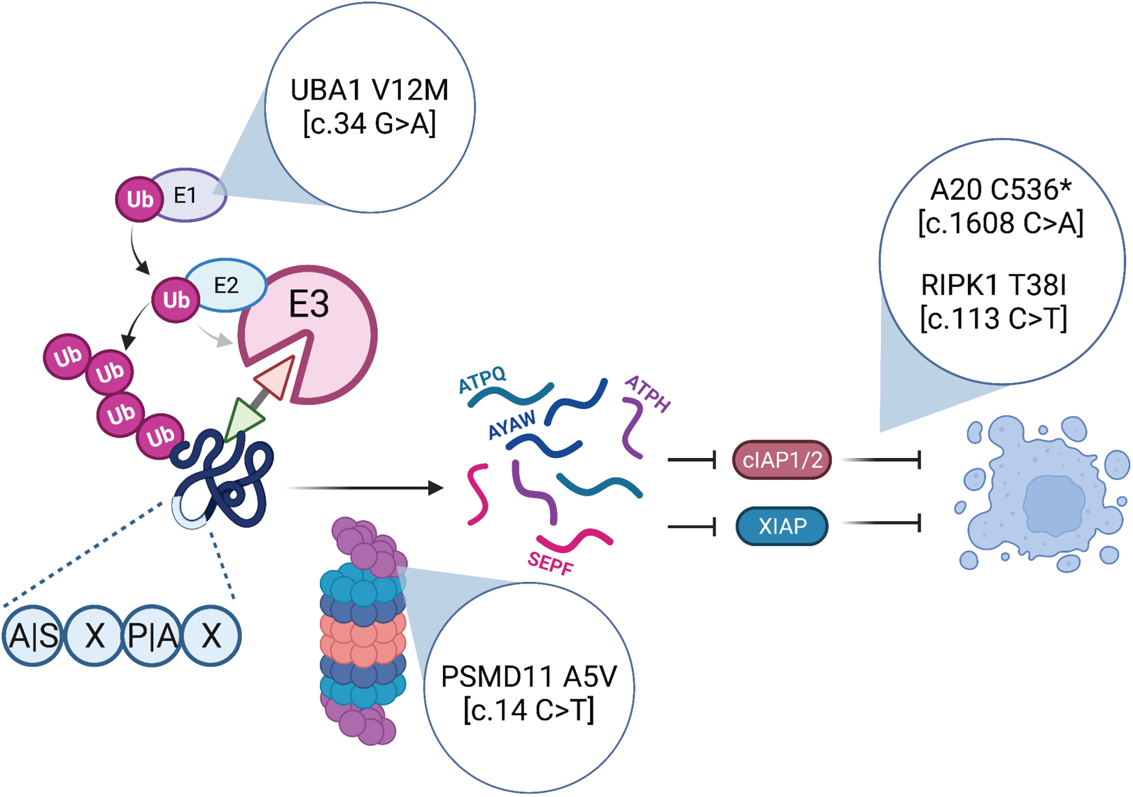
| Proposed model of MZ1 resistance in 22Rv1 cells. Summary of the whole-exome sequencing findings based on the identified protein-coding variants present specifically in the MZ1-resistant clones.

**Supplementary Fig. 5** contains all uncropped images of immunoblots and coomassie-stained gels in this manuscript.

## Supplementary Methods

### Whole Exome Sequencing processing and analysis

1E6 of 22Rv1 parental and resistant clones were collected and extracted for genomic DNA using Qiagen Blood & Tissue Kit according to the manufacturer’s manual. Purified genomic DNA was treated with 4 µL of RNase A (100 mg/mL) for 2 minutes at room temperature; RNase A was subsequently removed by QIAamp DNA Micro Kit following the Cleanup of Genomic DNA protocol. 0.5 µg of the DNA samples were library prepared and followed by exome capture and sequencing (Illumina). WES somatic variant discovery followed the workflow utilizing “Functional Equivalence” pre-processing with GATK4^61^. Paired end sequencing reads were mapped to the human reference genome (NCBI Assembly GRCh38) using bwa-mem (0.7.15-r1140) (https://arxiv.org/abs/1303.3997) with default settings. Variant calling was performed using Mutect2 (GATK 4.1.4.1)^62^, LoFreq2 (v2.1.3.1)^63^ and Strelka (v1.0.15)^64^ followed by functional annotations by Ensembl Variant Effect Predictor (release 99)^65^. Only variants that were identified by at least two of the three callers were selected for examination.

